# HIV-1 Mutants that Escape the Cytotoxic T-Lymphocytes are Defective in Viral DNA Integration

**DOI:** 10.1101/2021.08.29.458043

**Authors:** Muthukumar Balasubramaniam, Santosh Thapa, Benem-Orom Davids, Alex Bryer, Chaoyi Xu, Jiong Shi, Christopher Aiken, Jui Pandhare, Juan R. Perilla, Chandravanu Dash

## Abstract

ABSTRACT

HIV-1 replication is durably controlled in certain untreated HIV-1-infected individuals expressing particular human leukocyte antigens (HLA). These HLAs tag infected cells for elimination by presenting specific viral epitopes to CD8+ cytotoxic T-lymphocytes (CTL). In individuals expressing HLA-B27, CTLs primarily target the capsid protein (CA)-derived KK10 epitope. Selection of CA mutation R264K helps HIV-1 escape the CTL response but severely diminishes virus infectivity. Here we report that the R264K mutation-associated infectivity defect arises primarily from impaired viral DNA integration. Strikingly, selection of the compensatory CA mutation S173A or depletion of host cyclophilin A largely rescues the R264K-associated integration and infectivity defects. Collectively, our study reveals novel mechanistic insights into the fitness defect incurred by an HIV-1 variant escaping a CA-directed CTL response.

## INTRODUCTION

Over 35 million people have already died from the ongoing HIV/AIDS pandemic, and no curative therapy is available for the ∼38 million people currently living with the virus. The standard of care is antiretroviral therapy (ART) involving combination of drugs targeting HIV-1-encoded enzymes^1^. The ART, by controlling the viral load and preventing disease progression, has significantly reduced the mortality and morbidity associated with HIV-1 infections^2-6^. However, because ART does not eradicate the virus^7^, evolution and spread of drug-resistant virus strains significantly increase the risks of HIV-1 transmission and disease progression^8-12^. Therefore, continued research directed towards novel therapeutic^13-16^ and preventive strategies are urgently needed to mitigate the devastating impact of the global HIV pandemic.

There is strong evidence that the host immune system can naturally control HIV and delay disease progression^17-21^. For instance, the virus replication is durably suppressed and disease progression is delayed in a small subset of untreated HIV-1-infected individuals known as “elite controllers”^22-24^. Such long-term asymptomatic infection in the absence of ART has been attributed to robust immunological control mechanisms coordinated by human leukocyte antigens (HLA) ^25-28^ and CD8+ cytotoxic T-lymphocytes (CTLs)^29-37^. Notably, a strong association between HIV-1 control and presentation of viral epitopes by HLA-B57 or HLA-B27 to HIV-1-specific CTLs has been demonstrated^18, 26, 38-45^. However, CTL-mediated HIV-1 control is compromised by escape mutations that diminish recognition of viral epitopes by the HLA and/or CTLs^46-53^.

CTL response against the HLA-B27-restricted KK10 epitope^54^, derived from the capsid (CA) domain of the HIV-1 Gag protein, has been linked to robust virus control and delayed disease progression^17, 38, 55-57^. The immunological constraint on virus replication eventually leads to the emergence and selection of an HIV-1 variant containing mutations in the KK10 epitope of CA^55, 58-63^; specifically, the prerequisite mutation L268M (LM) emerging early followed by the consequential escape mutation R264K (RK) selected late in the course of infection^46, 55, 64^. The CTL escape mutation RK inhibits the binding and presentation of the CA-derived mutant peptide by HLA-B27^55, 65^ but also inflicts a significant fitness cost to the infectivity of the virus^46, 51, 55, 61, 64-67^. This is unsurprising because CA is genetically the most fragile of all HIV-1 proteins^68, 69^ and is a key player in multiple stages of viral replication^70-78^. For instance, the CA forms the conical capsid shell that houses the viral genome and other viral/cellular factors necessary for infection^79-85^. CA also provides critical binding interfaces for key host factors like cyclophilin A (CypA)^86-88^ and cleavage and polyadenylation-specific factor 6 (CPSF6)^89, 90^ to promote early steps of HIV-1 infection^91-98^. Remarkably, the infectivity defect of the RK mutant virus is largely restored upon selection of the compensatory CA mutation S173A (SA)^61^ or upon depletion of CypA^61, 66^. However, the mechanism(s) by which these CTL-escape CA mutations reduce or restore HIV-1 replication remains largely undefined.

Here we report the comprehensive biology of the CTL-escape RK mutant virus by using a multi-pronged approach that included structure-guided molecular dynamics (MD) simulations, binary protein interaction studies by yeast two hybrid (Y2H) assay, virus infectivity assays, biochemical characterization of isolated viral replication complexes, and genetic analysis of the viral genome. Collectively, our results have uncovered reduced HIV-1 integration as the principal mechanism underlying the infectivity defect of the RK variant. Notably, these findings also provide further evidence for a regulatory role of CA in post-nuclear entry steps of HIV-1 infection and reinforces the need of research focused on CA-targeting inhibitors and CTL-based vaccines.

## RESULTS

### CTL escape-associated CA mutations do not alter capsid structure

The HIV-1 capsid is assembled through specific intra- and inter-molecular interactions between ∼1500 copies of CA monomers organized into a lattice of ∼240 hexameric and 12 pentameric units^79, 80, 99-101^. Since the structural integrity of the capsid is essential for infection^102^, we employed structure-based molecular modeling and MD simulations to probe whether the CTL escape CA mutations alter stability of the CA hexamers and CA tubular assemblies. Fig. 1A depicts the relative positions of the CTL escape-associated amino acids [R264(R132), L268(L136), and S173A(S41)] and the substitution changes [R264K, L268M, S173A] linked to the HLA-B27-restricted KK10 epitope in the CA. All three amino acids—R264, L268, and S173—are located in the NTD-NTD interface between two neighboring CA monomers and are positioned on the exterior of the CA hexamer (Fig. 1B) and pentamers (Fig. 1C). Notably, none of the three amino acids is in the proximity of CypA binding loop (BL) or the CPSF6 binding pocket.

**Figure 1.**
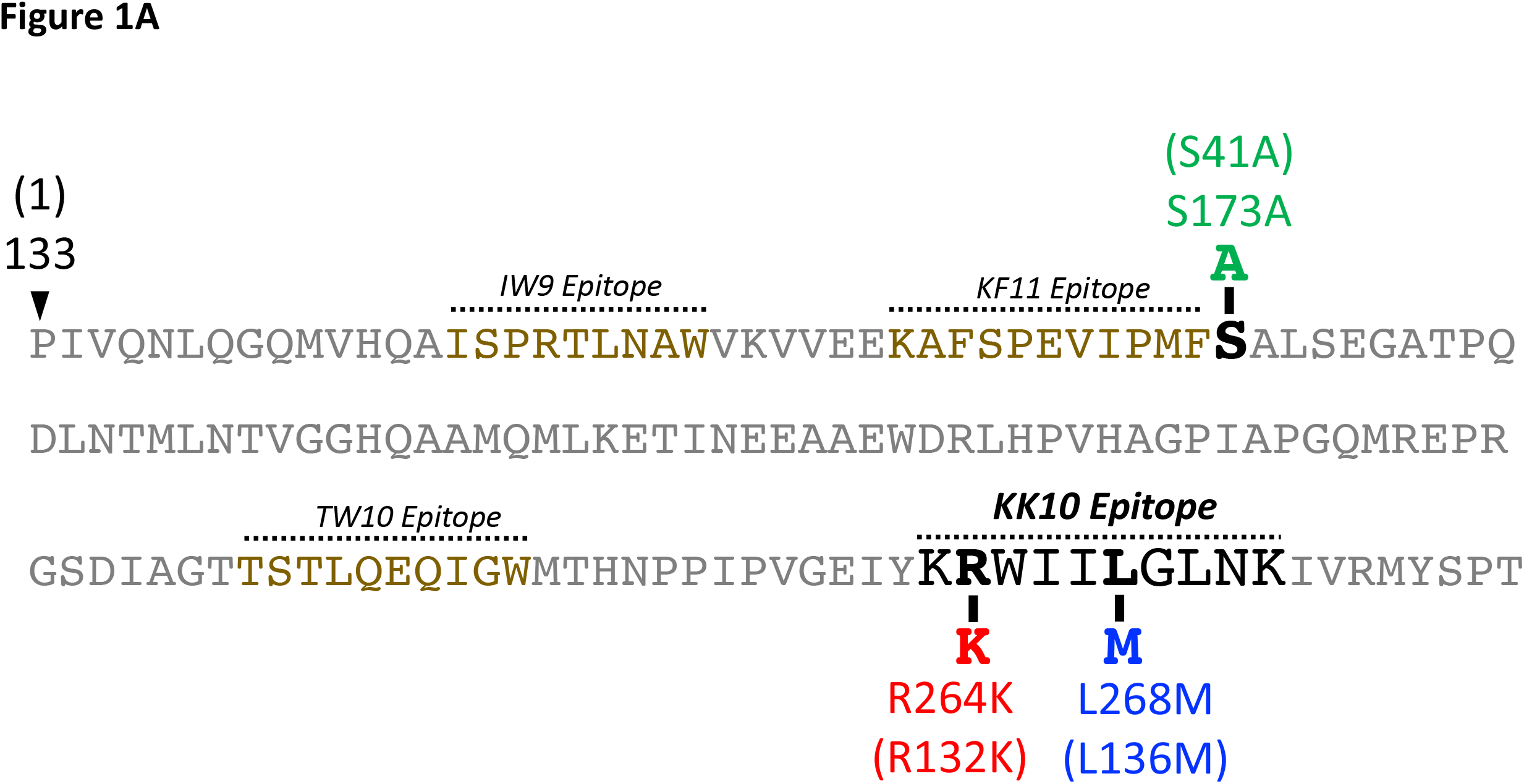
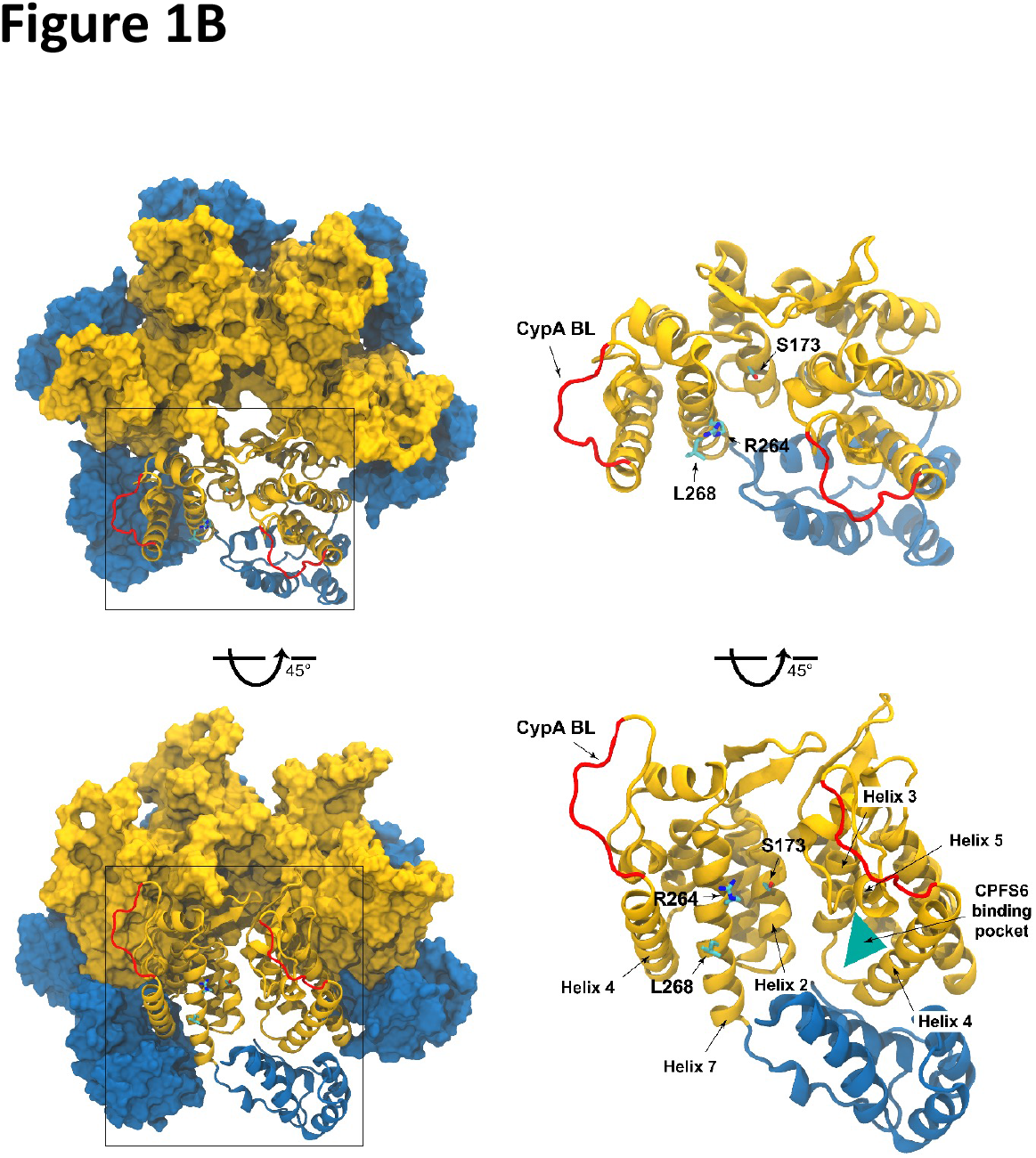
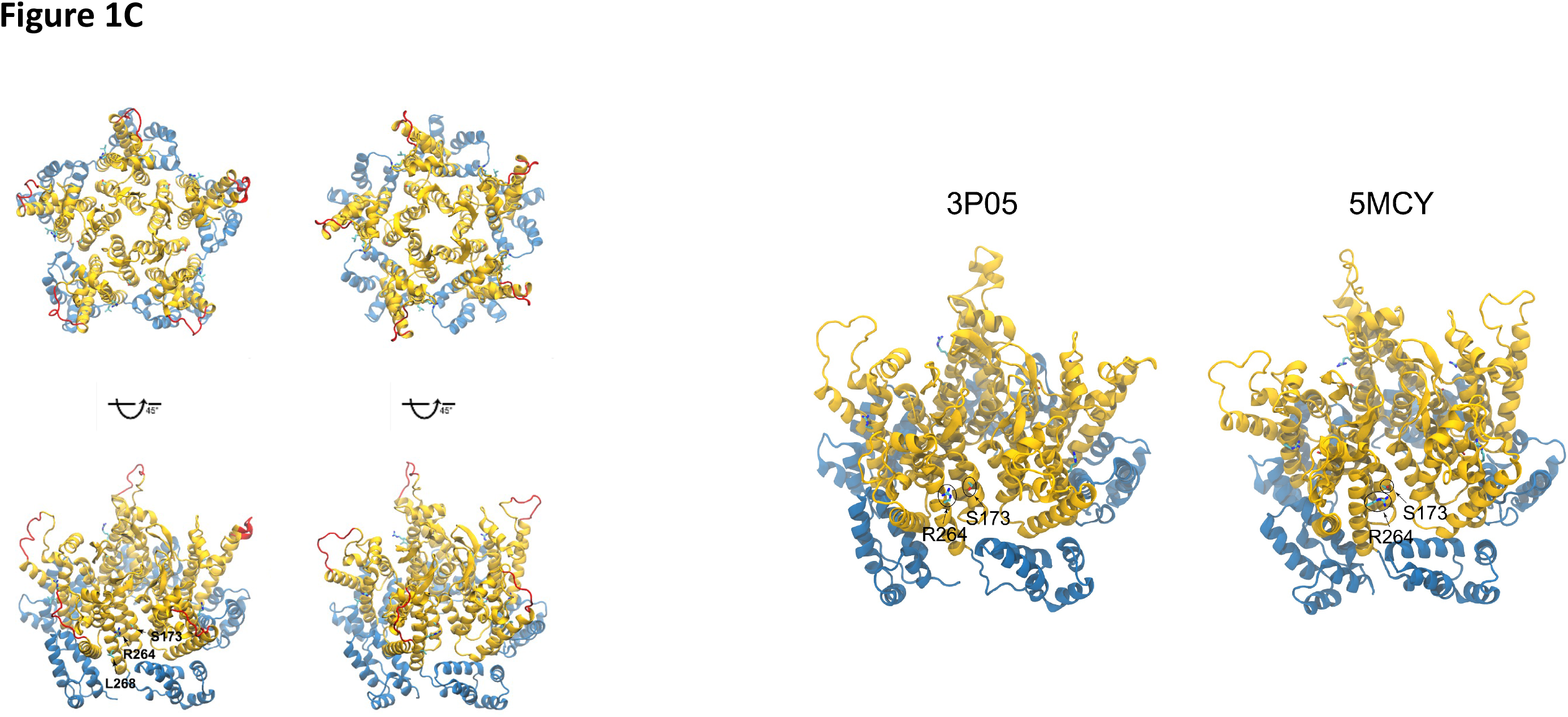
**(A)** Location of CTL-targeted epitopes in the HIV-1 capsid N-terminal domain and the CTL escape-associated amino acid changes linked to the HLA-B27-restricted KK10 epitope. The standard Gag numbering of the amino acids is followed, and the capsid numbering of the amino acids are in parentheses for reference. **(B)** Relative positions of three CA residues, R264(R132), L268(L136) and S173(S41) in the context of a WT CA monomer and hexamer. (Left) A full length native HIV-1 CA hexamer (PDBid: 4XFX) is represented by a surface model. The N-terminal domain (NTD) is colored in gold, while the C-terminal domain (CTD) is in blue. (Right) a CA monomer and the N-terminal domain from one of its neighboring monomers are highlighted in a ribbon representation. The positions of R264, L268 and S173 residues in the NTD-NTD interface are labelled, CypA binding loop (BL) is colored in red, and the CPSF6 binding pocket is depicted as a green triangle. **(C)** Relative positions of three CA residues, R264(R132), L268(L136) and S173(S41) in the context of a WT CA pentamer. (Left) Ribbon representation of a full-length native HIV-1 CA pentamer. The N-terminal domain (NTD) is colored in gold, while the C-terminal domain (CTD) is in blue. The positions of R264, L268 and S173 residues in the NTD-NTD interface are labelled and the CypA binding loop (BL) is colored in red. (Right) R264 and S173 residues on two CA pentamer models, PDB accession codes 3P05 and 5MCY.

MD simulations showed similar dynamic properties of the WT and CTL escape mutant hexamers (Fig. 2). Specifically, the Cα root mean square deviations (RMSDs) of all the CA hexamer systems showed an increase at 20 ns and were plateaued at around 2Å after 50 ns (Fig. 2A). Importantly, the mean Cα root mean square fluctuations (RMSFs) of all systems exhibited negligible differences, especially in the structured regions (Fig. 2A), and the structural flexibilities of all the hexamer systems were comparable (Fig. 2B). The inter-hexamer interactions of the CA residues R264, L268, and S173 in the WT hexamer are maintained in the mutant hexamers (Fig. 2C). For example, the top two contacts made by R264 (with residues I261 and I267), by L268 (with residues W265 and N271), and by S173 (with residues S176 and P170) are preserved in mutant hexamers. While the LM and SA substitutions preserve the electrostatic potential, the RK substitution renders the sidechain a relatively more positive potential (Fig. S1). Nevertheless, the overall electrostatic potentials of all the mutant hexamers are very similar to the WT hexamer (Fig. S1). Further, we probed for the presence of ions in the CA hexamers to better understand the interactions between the capsid and its native environment (Fig. S2). Specifically, chloride ions form an inner layer adjacent to the surface of the capsid, while sodium binds to the exterior of the capsid. Examination of the transfer rates for both ionic species present in the simulation reveal regions of high occupancy for chloride (cyan) and sodium (yellow) (Fig. S2). Comparison of ion occupancy of WT hexamers to that of the RK, RKLM and RKLMSA hexamers, showed no significant difference in the interaction between the capsid and the ionic environment.

**Figure 2.**
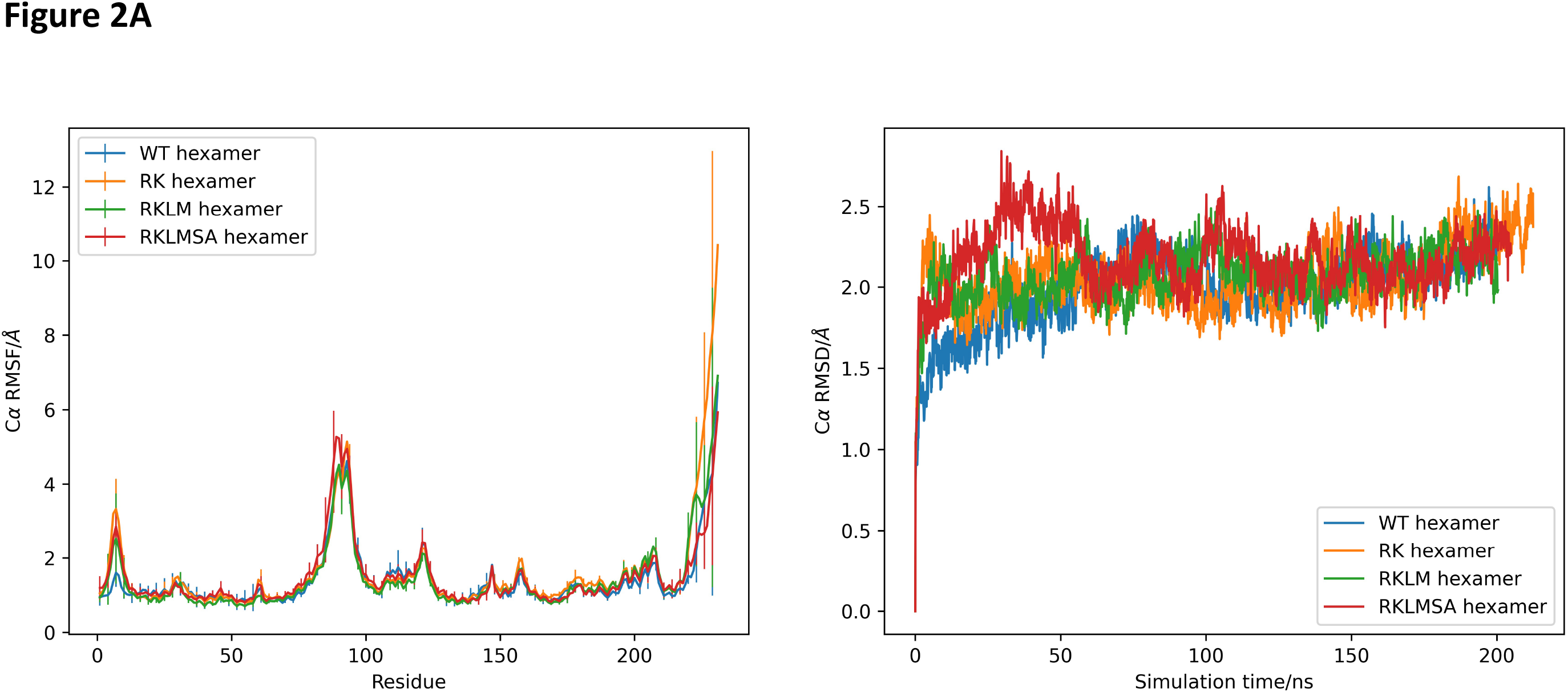
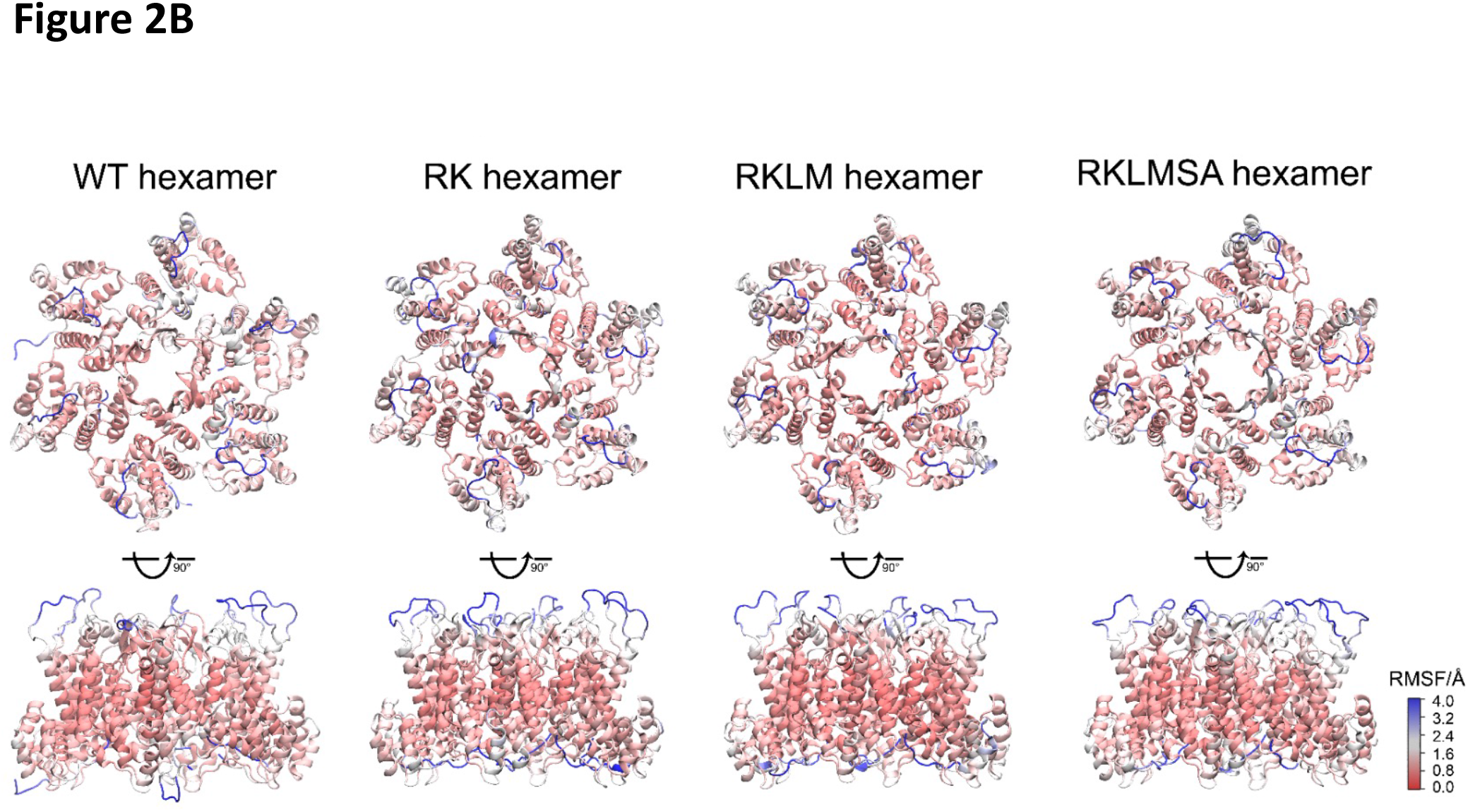
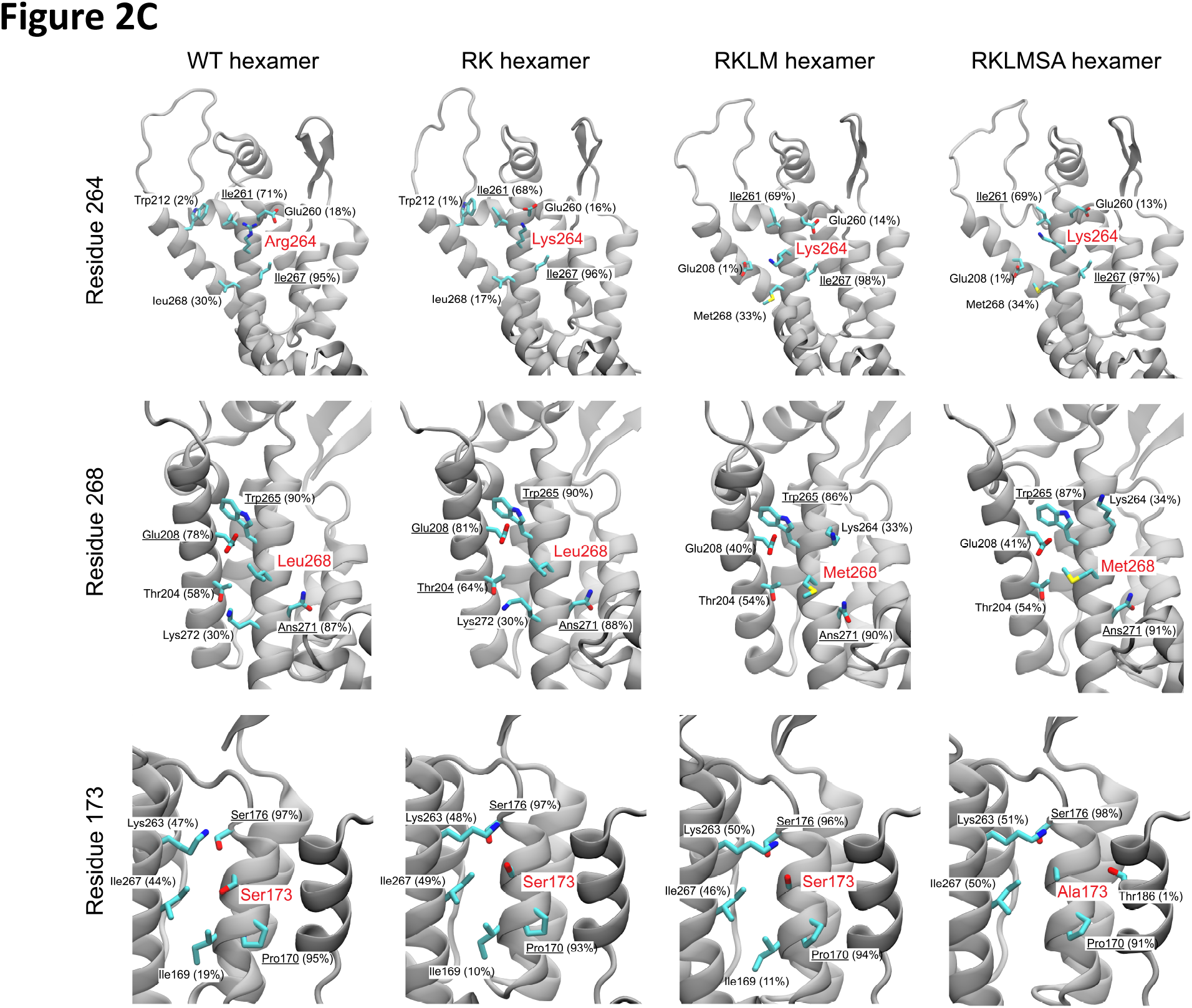
**(A)** Structural stability of HIV-1 CA hexamer influenced by three CA residue substitutions, R264(R132), L268(L136) and S173(S41). (Left) The average Cα root mean square fluctuations (RMSFs) of the HIV-1 WT, RK, RKLM, and RKLMSA hexamers derived from molecular dynamics simulations are shown. (Right) The Cα root mean square deviations (RMSDs) of HIV-1 WT hexamer and 3 mutation hexamers from molecular dynamics simulations. **(B)** Structural flexibilities of the WT and three mutation hexamers, colored by the average Cα RMSF values, increasing from red to blue. **(C)** Inter-hexamer interactions of three CA residues, R264(R132), L268(L136) and S173(S41), maintain after substitutions. Top 5 inter-hexamer contacts involved in CA residue 132 (first row), 136 (middle row) and 41 (last row) are indicated. The corresponding contact occupancies are in the parenthesis after residue names. The residue names with occupancies greater than 60% are underlined. The formation of a residue contact is defined as the distance between sidechains from not neighboring residues within 3.0 Å.

Furthermore, we constructed and simulated RK, RKLM and RKLMSA mutant CA tubes to test the effects of these mutations on the structure and dynamics of the tubular capsid assembly (Fig. 3). After 20 ns of MD equilibration, no structural differences between any of the CA tubes were observed. Tube radii and length values vary by no more than 0.1-0.2 nm for all mutant systems studied (Fig. 3A). Moreover, the conformational distributions of each mutation site were also quite similar (Fig. 3B), as well as the RMSF of capsid residues (Fig. 3C), indicating that mutations do not alter the dynamics of capsid lattice. These observations indicate the structural and dynamic similarity between these mutant CA tubes despite the presence of mutations.

**Figure 3.**
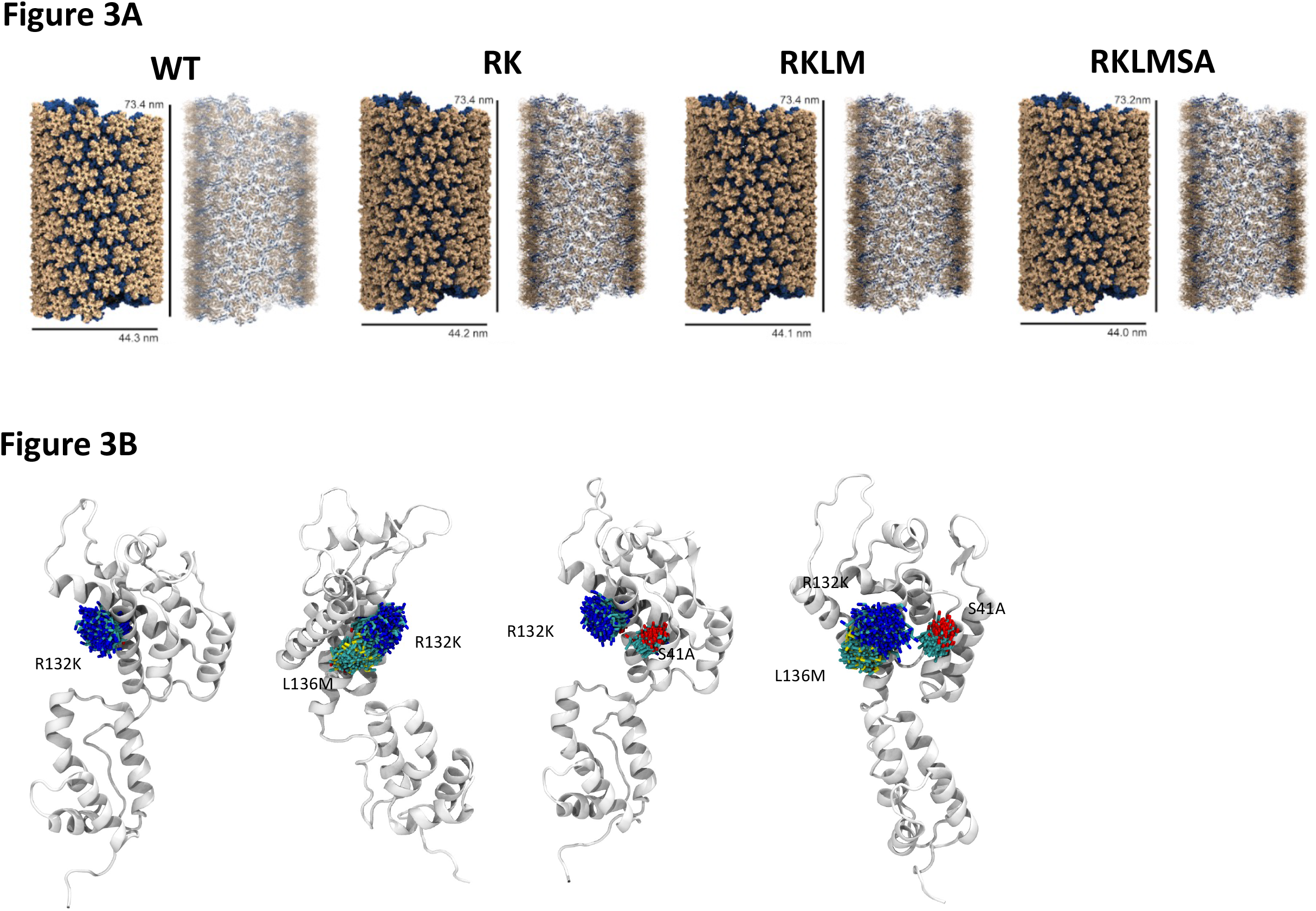
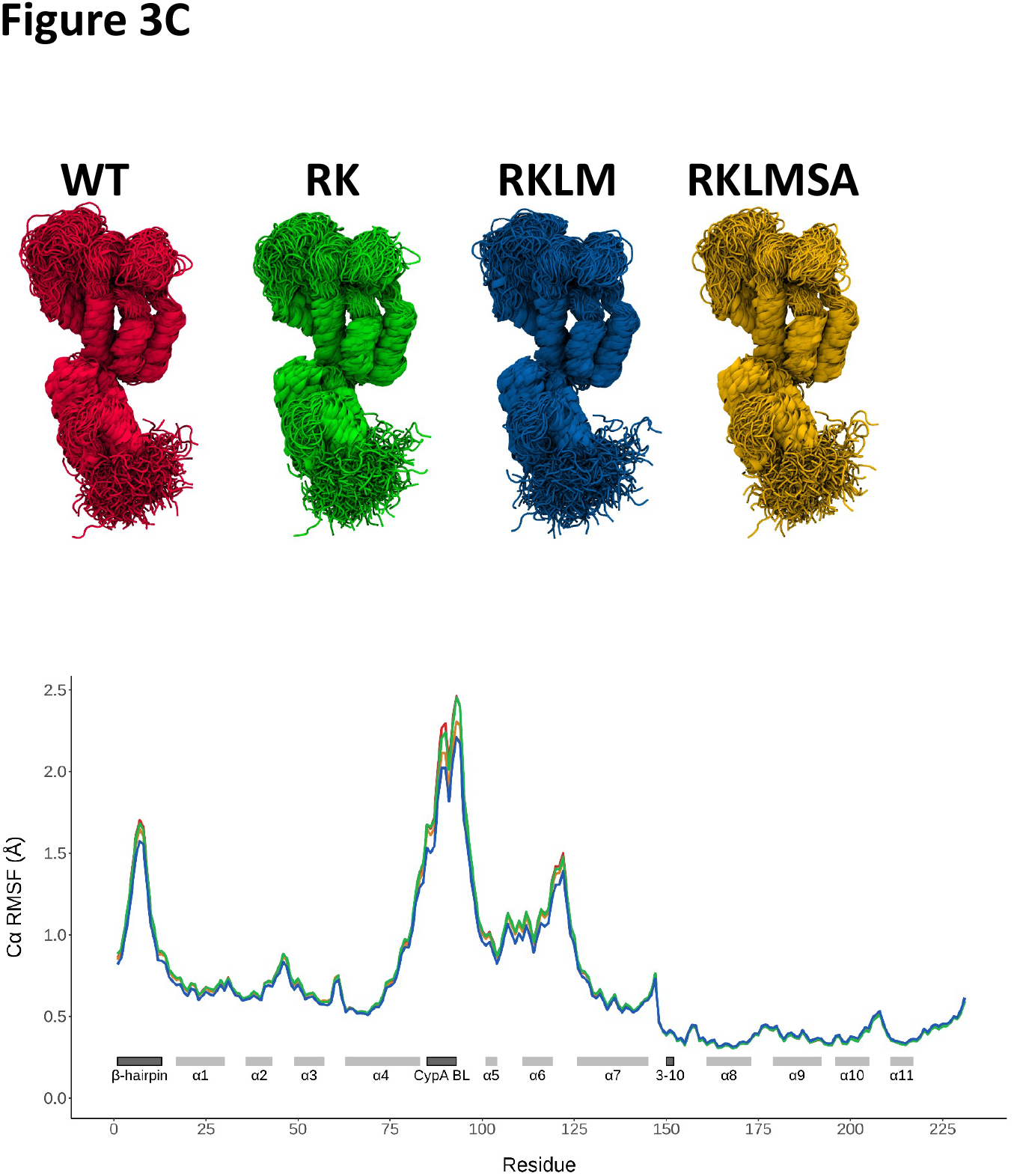
**(A)** Effects of the CTL-escape mutation on structure of CA tube assembly. **(B)** Effects of the CTL-escape mutation on conformational distribution of CA monomers. Each mutation site, in each system, aligned to a single monomer, demonstrating the conformational distributions of these residues. **(C)** Effects of the CTL-escape mutation on the root-mean-square fluctuation (RMSF) of capsid residues. (Top Panel) Each monomer is shown aligned by backbone regions in helices in the top row, colored accordingly. (Bottom) RMSF of Carbon alpha atoms in each CA tube system. RMSF values are averaged for all monomers.

The HIV-1 capsid has many binding interfaces for cellular co-factor recognitions ^72, 103^. For example, CypA selectively bridges two neighboring CA hexamers using canonical and non-canonical binding sites^88, 104^, both located at the CypA binding loop (BL). The CPSF6^90^ and Nucleoporin 153 (NUP153)^105^ target the CA NTD-NTD interface. Other co-factor binding sites on capsid include the center pore of CA hexamer [to recognize IP6^106^, nucleotide^107^, and Fasciculation and Elongation Protein Zeta 1 (FEZ1)^108^] and the tri-hexamer region [to recognize MX dynamin like GTPase 2 (MXB)^109^]. However, the locations of the CTL escape RK mutations are far away from these co-factor binding sites on the capsid (Fig. 1B) and render almost identical structures and dynamics of the CA hexamer and tube, which are unlikely to affect these co-factor recognitions. Together, our *in silico* studies indicate that the CTL escape-associated CA amino acid changes linked to the HLA-B27-restricted KK10 epitope do not alter the structural integrity and dynamics of the CA hexamers and capsid lattice, as well as the capsid-cofactor interactions.

### CA mutation RK does not perturb CA-CA and CA-host protein interactions

We next assessed the effect of the RK mutation on CA-CA and CA-host factor interactions by yeast GAL4-based two-hybrid (Y2H) assay^110-112^. We focused on the RK mutation because it is the primary determinant of the infectivity defect associated with HLA-B27 CTL escape ^61^. First, we examined the well-defined dimerization of WT CA monomers. The yeast cells co-transformed with the plasmid pairs encoding the GAL4 DNA Binding Domain (BD) in fusion with the WT CA (BD-CAWT) and the GAL4 Activation Domain (AD) in fusion with the WT CA (AD-CAWT), exhibited histidine prototrophy and *α*-galactosidase activity (Fig. 4A, Top panel), thus indicating direct protein-protein interaction. Yeast cells co-transformed with plasmid pairs encoding BD-CAWT and AD-mCherry or AD-CAWT and BD-mCherry fusion proteins exhibited auxotrophic growth on selection media lacking leucine and tryptophan but did not exhibit histidine prototrophy and *α*- galactosidase activity. These results highlight specificity of the Y2H assay in detecting bona fide CA-associated binary protein interactions.

**Figure 4.**
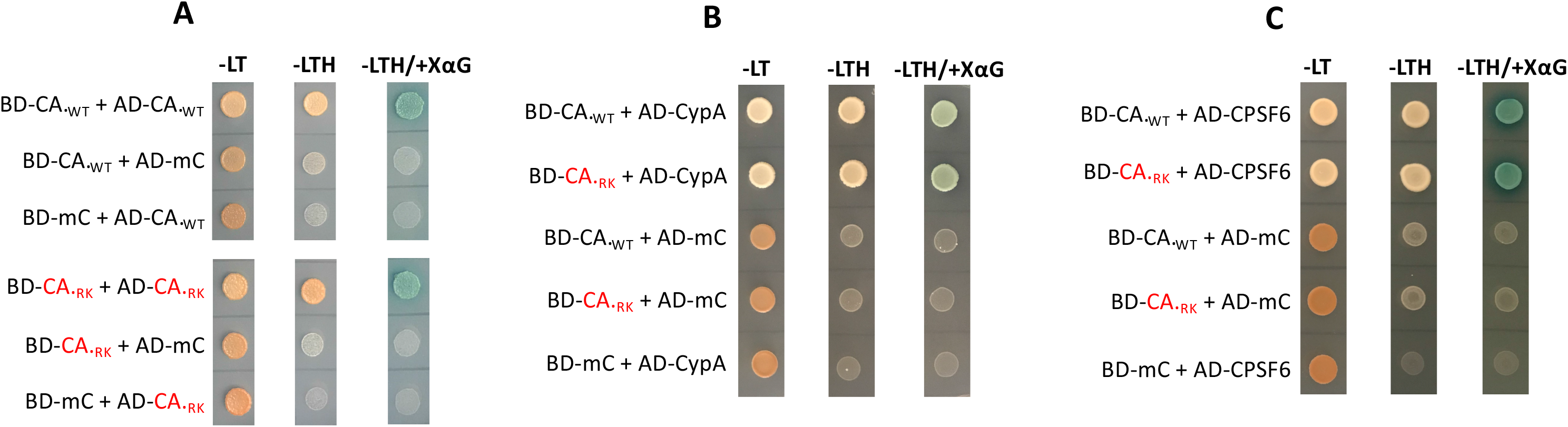
Effect of KK10-linked CTL escape mutation R264K on CA interactions. The effect of RK mutation on **(A)** CA:CA, **(B)** CA:CypA, and **(C)** CA:CPSF6 interactions were assessed by using GAL4-based yeast two-hybrid assays as described in Materials and Methods. Equal number of yeast cells co-transformed with the plasmid pairs encoding the GAL4 DNA Binding Domain (BD) in fusion with the bait protein and the GAL4 Activation Domain (AD) in fusion with the prey protein were spotted on synthetic dropout (SD) agar plates lacking leucine and tryptophan (-LT) or leucine, tryptophan, and histidine (-LTH) or leucine, tryptophan, and histidine but supplemented with X-*α*-D-galactoside (-LTH/+X*α*G) and were incubated at 30°C to assess for histidine prototrophy and *α*-galactosidase activity. (A-Top Panel) Growth of yeast co-expressing BD-CAWT and AD-CAWT, BD-CAWT and AD-mCherry (AD-mC), AD-CAWT and BD-mC on selection media. (A-Bottom Panel) Growth of yeast co-expressing BD-CARK and AD-CARK, BD-CARK and AD-mC, and AD-CARK and BD-mC on selection media. **(B)** Growth of yeast co-expressing BD- CAWT and AD-CypA, BD-CARK and AD-CypA, BD-CAWT and AD-mC, BD-CARK and AD-mC, and BD-mC and AD-CypA on selection media. **(C)** Growth of yeast co-expressing BD-CAWT and AD- CPSF6, BD-CARK and AD-CPSF6, BD-CAWT and AD-mC, BD-CARK and AD-mC, and BD-mC and AD-CPSF6 on selection media.

Importantly, we observed histidine prototrophy and *α*-galactosidase activity by the yeast co-expressing BD-CARK (BD in fusion with the RK CA) and AD-CARK (AD in fusion with the RK CA), but not by the yeast co-expressing BD-CARK and AD-mCherry or AD-CARK and BD-mCherry (Fig. 4A, Bottom panel). These results demonstrate that the CTL-escape RK mutation does not disrupt CA-CA binary interaction. Next, we tested whether the RK mutation affected interaction of CA with two key host factors-CypA and CPSF6 that facilitate post-entry steps of infection and are implicated in the infectivity defect of the RK variant virus. Interestingly, histidine prototrophy and *α*-galactosidase activity was observed by the yeast co-expressing BD-CAWT or BD-CARK and AD- CypA (Fig. 4B) or AD-CPSF6 (Fig. 4C), but not by the yeast co-expressing one of these fusion proteins and the complementary negative control mCherry fusion protein. These results show that the RK mutation has no significant effect on the binary interactions of CA with either CypA or CPSF6. Collectively, the Y2H assay data strengthen our *in silico* studies and indicate that the CTL escape CA mutation RK does not compromise the structural integrity of CA or its functional interactions with key host factors.

### RK mutation impairs HIV-1 infection but the compensatory mutation SA restores infectivity

HIV-1 escape from the KK10 epitope-targeted CTL response is functionally linked to specific CA mutations^55, 58, 59, 61, 62^. The prerequisite LM mutation, independently, has no significant effect on HLA binding, CTL recognition, and viral infectivity^61, 113^. However, the RK mutation independently or in combination with LM (RKLM) significantly reduces the binding of the KK10 epitope to the HLA-B27 molecule and drastically impairs infectivity^61-63, 65^. Strikingly, the compensatory CA mutation S173A confers WT-level, or more, infectivity to the RK and RKLM mutant viruses^61, 67^. Therefore, to understand the precise mechanism by which these CTL-escape CA mutations affect HIV-1 infectivity, we compared the infectivity of the WT and mutants using replication-competent virus particles. The LM mutation alone does not significantly affect infectivity and hence was not included in the assessment.

For the TZM-bl-based infectivity assay, we inoculated TZM-bl cells^114^ with equivalent MOI of WT or the mutant virus and 48 hours post infection (hpi) assessed the luciferase activity in the cellular lysates. Compared to the WT virus, the infectivity of the RK and the RKLM mutant viruses were drastically reduced, by up to 10-fold (Fig. 5A). The comparable infectivity of the RK and the RKLM variants indicates that the RK, and not the LM mutation, is the principal determinant of the infectivity defect. Importantly, the addition of the compensatory SA mutation to the RKLM virus led to restoration of infectivity to the level of the WT virus (Fig. 5A and 5B). For the infectivity assays using Jurkat T-cells, we spinoculated cells with equivalent MOI of WT or any of the mutant viruses and assessed the HIV-1 Gag protein levels by flow cytometry. As shown in Fig. 5C-D, in comparison to the WT virus, the infectivity of the RK and RKLM mutant viruses were significantly reduced and the compensatory SA mutation led to significant restoration of its infectivity. Collectively, our infection assays in two different cell types establish that the CTL-escape CA mutation RK severely impairs HIV-1 infectivity and this defect is mostly rescued by the compensatory SA mutation.

**Figure 5.**
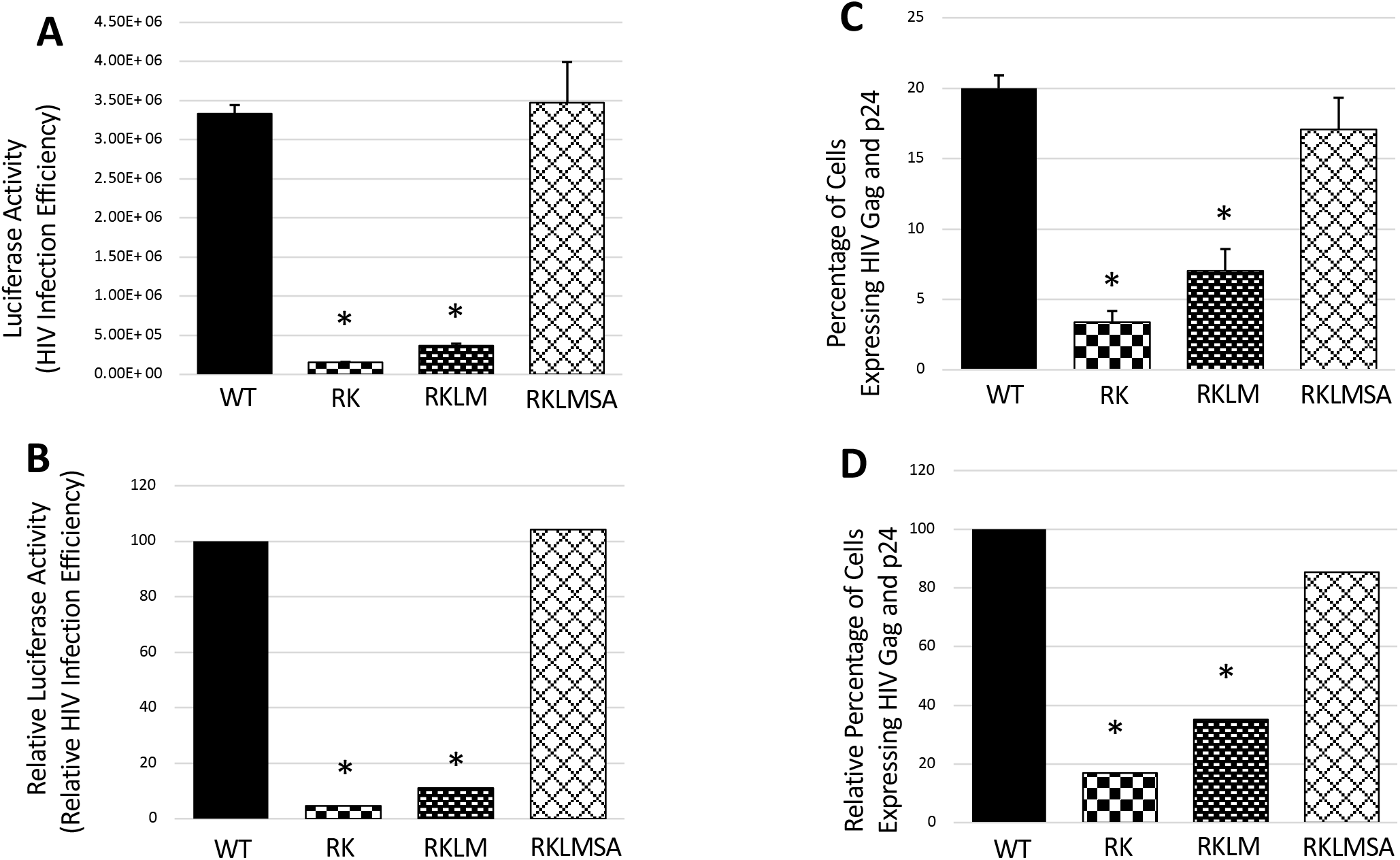
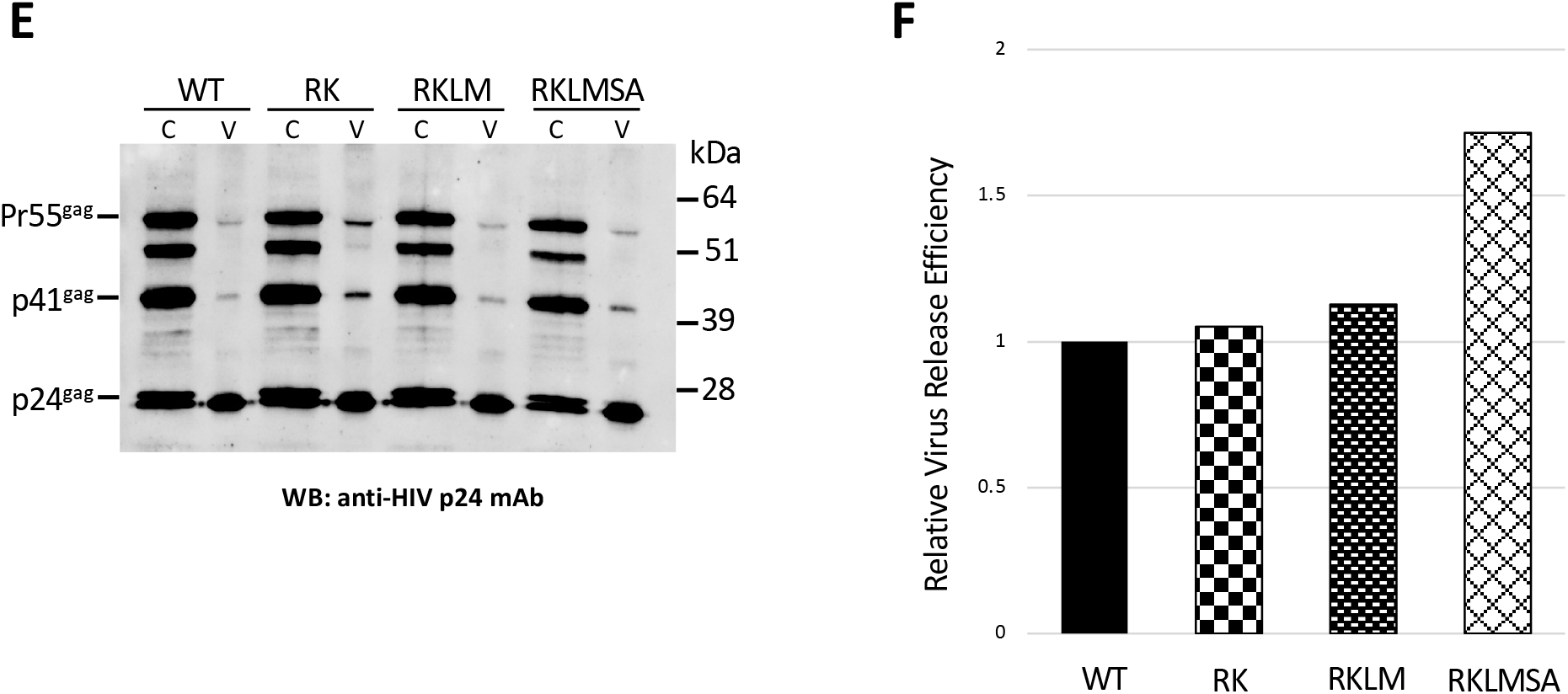
Effect of KK10-linked CTL escape mutations on HIV-1 infectivity. Infectivity of WT and mutant viruses were determined using the **(A and B)** TZM-bl indicator cell line and **(C and D)** Jurkat T-cells as described in Materials and Methods. TZM-bl cells inoculated, in replicates, with the WT or mutant viruses for 2 hours were further cultured for 48 hours and then lysed. **(A)** Luciferase activity in the cell lysates, indicative of virus infectivity, was measured as luminescence and, after subtracting the luciferase activity in mock infected cells from the data, were plotted as relative light units (RLU). **(B)** Relative luciferase activity, indicative of percentage infection efficiency, was plotted relative to WT. Jurkat cells spinoculated, in replicates, with the WT or mutant viruses for 2 hours at 37°C were further cultured for 48 hours at 37°C, after which the cells were fixed and stained with FITC-labelled antibody that identifies viral Gag and p24. **(C)** The percentage of FITC-labelled cells (marker for HIV-1 infectivity) was determined using FACS analysis. **(D)** Relative Gag and p24 expression, indicative of percentage infection efficiency, was plotted relative to WT. Data shown are representative of at least three independent experiments, with error bars representing the SEMs. Effect of KK10-linked CTL escape mutations on late stages of HIV-1 replication. HEK293T cells transfected with WT or the mutant proviral clones were cultured for 48 hours at 37°C and the viral particles in the culture media was collected by ultracentrifugation. **(E)** Cell (C) and virus (V) lysates were resolved by SDS-PAGE and the virus proteins were probed by western blotting. **(F)** The virus release efficiency was determined as described in Materials and Methods.

### RK-associated infectivity defect is not due to block in the late stages of virus replication

HIV-1 CA plays key roles during early and late events of virus replication. During the late stages, the CA-CA interaction is essential for hexameric Gag lattice formation and assembly of immature progeny viruses^115, 116^. Furthermore, cleavage of CA from the Gag precursor is critical for the formation of the conical capsid of the mature virions^117^. Therefore, the impaired infectivity of the RK mutant virus (Fig. 5A-D) could be a consequence of the CA mutation adversely affecting any of these late-stage processes. To test this possibility, we probed the effects of the CTL escape CA mutations on Gag processing and virus particle production.

HEK293T cells were transfected with the molecular clones of the WT or the CA mutants and cultured for 48 hours, after which the cells and the corresponding culture supernatants were collected. The virus proteins from the cells and from the pelleted virus particles were probed by western blotting (Fig. 5E). The Gag processing efficiency in the producer cell was not significantly altered by any of the CA mutations. Further, none of the CA mutations were found to reduce virus release efficiency—measured as the ratio of the amount of virus-associated CA to the total Gag (total amount of virus-associated CA, cell-associated Pr55Gag, cell-associated p41Gag and cell-associated CA) (Fig. 5F). Taken together, these results suggest that the CTL escape CA mutations do not significantly alter the late stages of HIV-1 replication and that the RK mutation-associated infectivity defect maps to one or more of the post-entry steps of the virus replication.

### RK mutation has no significant effect on HIV-1 reverse transcription and nuclear entry

The CA mutations RK and RKLM are not positioned to significantly alter the structural integrity of the capsid structures and, by extension, the reverse transcription of the mutant viruses. Intriguingly, the infectivity defect imposed by the RK and RKLM mutations have previously been reported to arise from reduction in reverse transcription^61^. So, we quantified levels of reverse transcription in T-cells inoculated with these viruses. Jurkat or SupT1 cells spinoculated^118^ for 2 hours with equivalent MOI of the WT or the mutant viruses were cultured for 24 hours, after which total DNA was isolated from the cell samples. The viral late RT products (i.e., viral DNA) in the samples was measured by qPCR and the copy numbers of the viral DNA were calculated by interpolation from a standard curve. The results revealed that the RK and the RKLM mutations did not impose significant alterations in the reverse transcription levels (Fig. 6A-B) that could be correlated with the magnitude of the impairment in virus infectivity (Fig. 5A-D). The addition of the compensatory SA mutation to the RKLM virus had no significant effect on the reverse transcription levels, as well. These results diverge from published work^61^ and suggest that the RK mutation-associated infectivity defect arises from a block downstream of the reverse transcription step.

**Figure 6.**
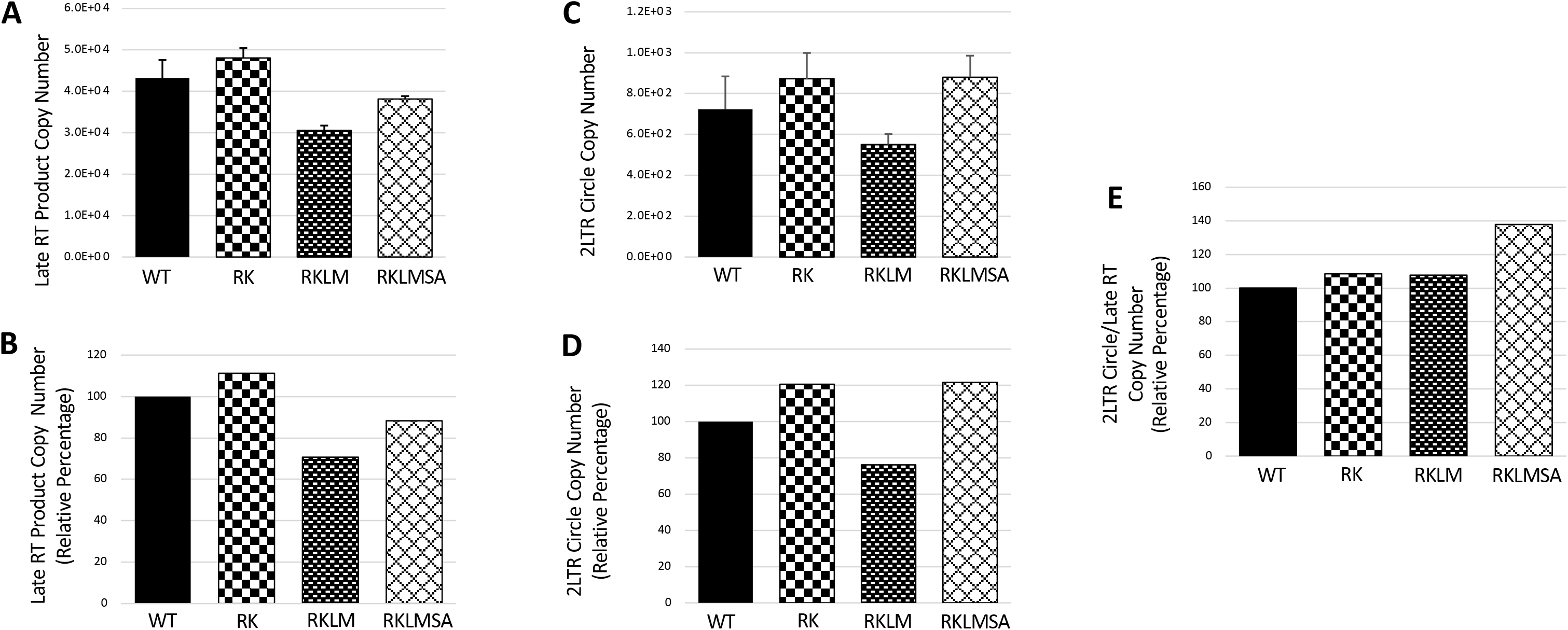
Effect of KK10-linked CTL escape mutations on HIV-1 reverse transcription levels and nuclear import efficiency. Copy number of late RT products and 2-LTR circles present in the total DNA isolated from Jurkat cells spinoculated with the WT or mutant viruses for 2 hours at 25°C and cultured for 24 hours at 37°C were determined as described in Materials and Methods. **(A)** Copy number of late RT products of WT or the mutant viruses as measured by qPCR and calculated using a standard graph of known copy numbers of HIV-1 molecular clone (surrogate marker for RT product). **(B)** Percentage late RT products plotted relative to WT. **(C)** Copy number of 2-LTR circles as measured by qPCR and calculated using a standard graph of known copy numbers of 2-LTR-containing plasmid DNA. **(D)** Percentage copy number of the 2-LTR circles plotted relative to WT. **(E)** Calculated ratio of the copy number of 2-LTR circles to the respective late RT products plotted as percentage relative to WT. Data representative of three independent experiments, with error bars representing the SEMs.

HIV-1 nuclear import is the next step in the virus replication cycle and prior reports did not show the effect of RK mutation on viral nuclear import^61, 67^. So, we tested the nuclear import efficiency of the WT and the CA mutant viruses by quantitating the viral 2-LTR circles in infected cells. A canonical 2-LTR circle is generated in the nucleus when the two LTRs in an unintegrated viral DNA are joined by the host non-homologous DNA end-joining pathway^119, 120^. Therefore, the viral 2-LTR circles are the most commonly used surrogate marker for estimating the efficiency of the nuclear import of the viral DNA^121, 122^. The 2-LTR circles were measured by qPCR and their copy numbers were calculated by interpolation from a standard curve generated in parallel using known copy numbers of a plasmid containing a cloned 2-LTR sequence (Fig. 6C-D). The results show that the RK and the RKLM mutations did not impose significant reductions in the levels of nuclear import that could be correlated with the magnitude of the infectivity defect (Fig. 5A-D). Further, the RKLMSA virus did not exhibit any significant difference in the nuclear import levels. As the 2-LTR circles are derivatives of the late RT products and the RKLM virus showed a marginal reduction in reverse transcription (Fig. 6A-B) and nuclear entry (Fig. 6C-D), we calculated the ratio of 2-LTR circles to the late RT products to measure nuclear import efficiency (Fig. 6E). No significant difference in the nuclear import efficiency of the WT and CA mutant viruses was observed. These findings suggest that the CTL-associated CA mutations minimally affect the efficiency of reverse transcription and nuclear import of HIV-1.

### RK mutation blocks proviral integration and SA compensatory mutation restores it

Our results in Fig. 6 indicated that a post nuclear import defect is most likely responsible for the infectivity defect caused by the RK mutation. To test this, we used the *Alu*-gag nested PCR assay to compare the levels of chromosomally integrated viral DNA (proviral DNA) in cells inoculated with WT or the mutant viruses. The *Alu*-gag nested PCR assay takes advantage of the widespread occurrence of the *Alu* sequence in the human genome^123, 124^, the reported preferential integration of HIV-1 near such *Alu* sequences^125-127^, and the selective exponential amplification of only the chromosomally integrated viral DNAs in the 1^st^ round end-point PCR followed by their quantitation in the 2^nd^ round qPCR^128, 129^.

Jurkat or SupT1 cells spinoculated for 2 hours with equivalent MOI of the WT or any of the mutant viruses were cultured for 24 hours, after which total DNA was isolated from the cell samples. The copy numbers of proviral DNA were calculated by interpolation from a standard curve generated in parallel using known copy numbers of pNL43 during the 2^nd^ round qPCR (Fig. 7A-B). As the late RT products constitute the sole source of the proviral DNAs, the viral late RT products in the same samples were also measured by qPCR in parallel (Fig. 7C-D). The integration efficiency was calculated by plotting the data as the ratio of the copy numbers of proviral DNAs to the late RT products (Fig. 7E). Strikingly, the results revealed a significant reduction in the proviral DNA levels (Fig. 7A-B) and integration efficiency (Fig. 7E) of the RK and RKLM viruses when compared to the WT infection. Importantly, the addition of the compensatory SA mutation to the RKLM virus (RKLMSA) led to significant restoration of its integration efficiency (Fig. 7E), which largely correlated with the magnitude of the restoration of the virus infectivity (Fig. 5A-D). These findings, together with our data of reverse transcription and nuclear import, establish that the infectivity defect of the RK and the RKLM viruses are primarily due to the reduction in viral DNA integration into the chromosomal DNA.

**Figure 7.**
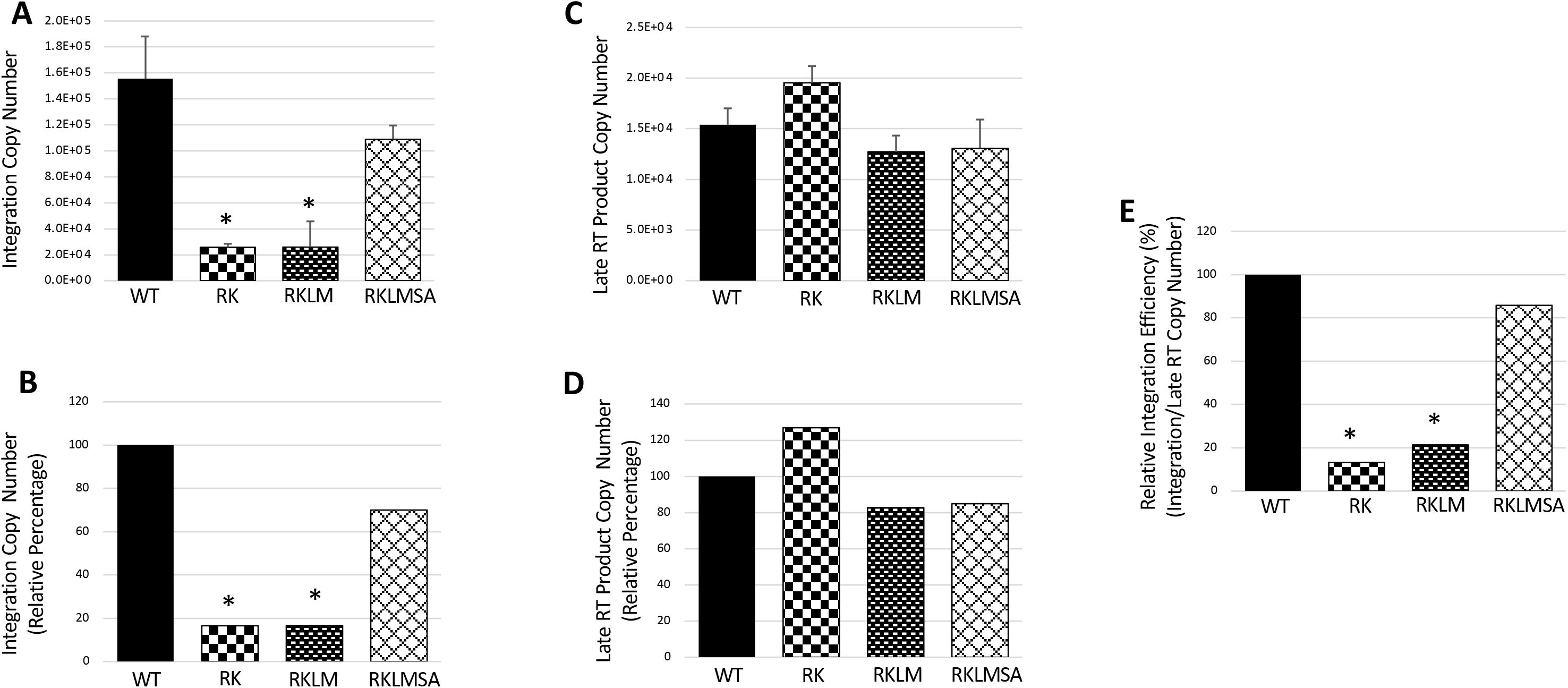
Effect of KK10-linked CTL escape mutations on HIV-1 proviral integration. Copy number of chromosome-integrated viral DNAs and late RT products present in the total DNA isolated from SupT1 cells spinoculated with the WT or mutant viruses for 2 hours at 25°C and cultured for 24 hours at 37°C were determined as described in Materials and Methods. **(A)** Copy number of chromosome-integrated viral DNAs as measured by *Alu*-Gag nested PCR and calculated by using a standard graph of known copy numbers of HIV-1 molecular clone (surrogate marker for viral DNA). **(B)** Percentage copy number of chromosome-integrated viral DNA plotted relative to WT. **(B)** Copy number of late RT products of WT or the mutant viruses as measured by qPCR and calculated using a standard graph of known copy numbers of HIV-1 molecular clone. **(D)** Percentage late RT products plotted relative to WT. **(E)** Calculated ratio of the copy number of chromosome-integrated viral DNAs to the respective late RT products plotted as percentage relative to WT. Data representative of three independent experiments, with error bars representing the SEMs.

### RK mutation has no significant effect on the integration activity of the cytoplasmic PICs

Integration of the HIV-1 DNA is carried out by the pre-integration complex (PIC)^130^. Therefore, the significant reduction in proviral integration of the RK variant could be the consequence of a reduction in PIC-associated viral DNA integration activity. To test this, we compared the integration activity *in vitro* of PICs extracted from cells inoculated with the WT or RK virus. Due to the well-recognized technical challenges associated with isolating the nuclear PICs, we used the cytosolic extracts from Jurkat or SupT1 cells^74, 131, 132^. The integration activity of these PICs is the measure of the number of viral DNA copies integrated into a heterologous target DNA and is quantified via a nested PCR assay. Intriguingly, compared to the WT PICs, there was no reduction in the integration activity of the RK mutant PICs (Fig. 8A-B and Fig. S3A-B). Notably, there was no reduction in the viral DNA content of the RK mutant PICs when compared to the WT PICs (Fig. 8C-D and Fig. S3C-D). Accordingly, the integration efficiency (i.e., the ratio of the integration activity to the viral DNA content) between the WT and the RK mutant PICs did not differ significantly (Fig 8E) or were comparable (Fig. S3E). Collectively, these results suggest that the integration activity of the RK mutant PICs is not defective and the RK mutation-associated reduction in proviral integration is not due to faulty assembly of PICs.

**Figure 8.**
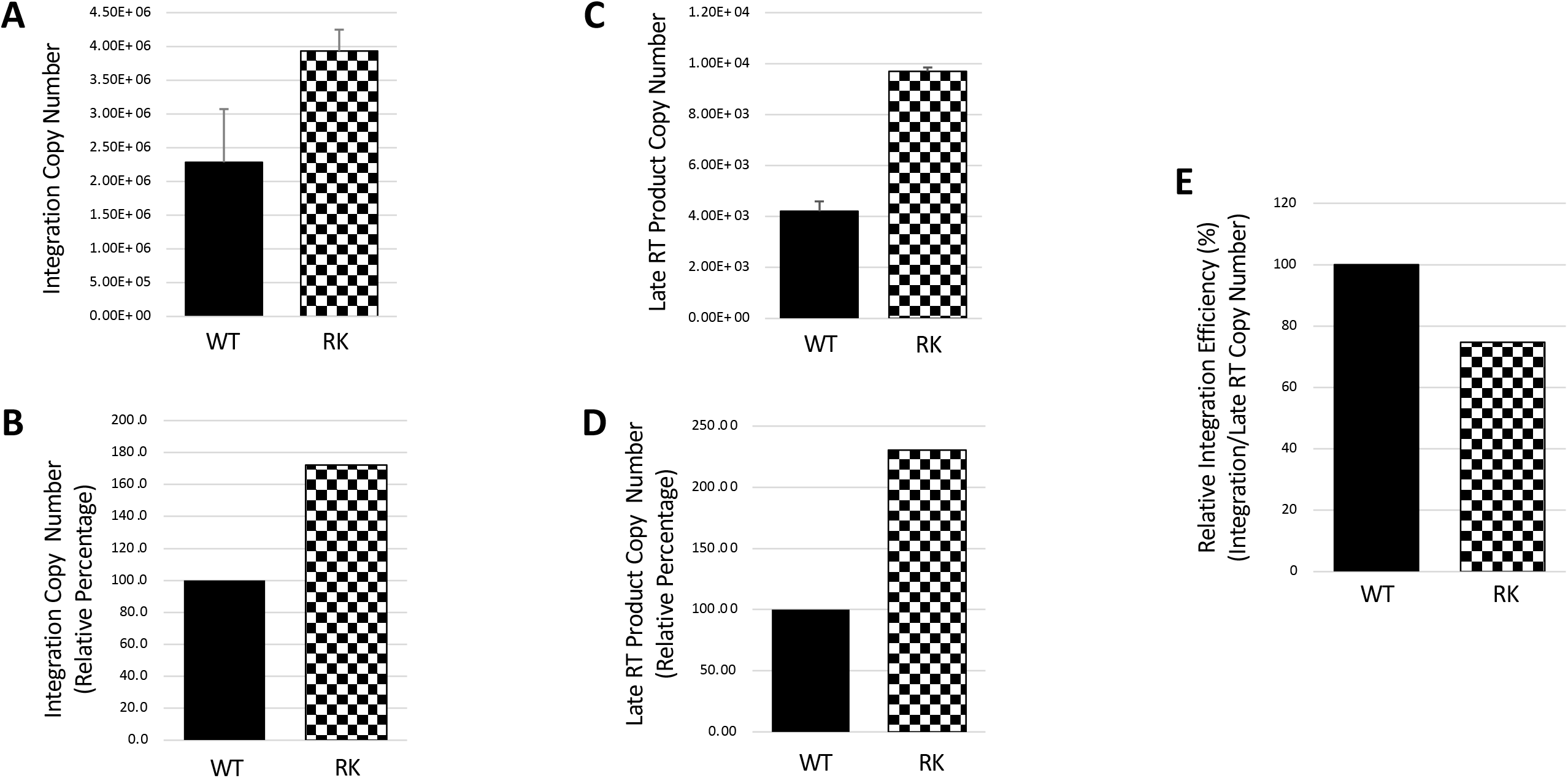
Effect of KK10-linked CTL escape mutation R264K on the integration activity in vitro of cytoplasmic PICs. Integration activity in vitro and the viral DNA content of the WT and RK cytoplasmic PICs isolated from Jurkat cells spinoculated with the respective viruses for 2 hours at 25°C and cultured for 5 hours at 37°C were determined as described in Materials and Methods. **(A)** Integration activity of the cytoplasmic PICs, i.e., the copy number of viral DNAs that were integrated into the target DNA during the in vitro integration assay, as measured by qPCR and calculated using a standard graph of known copy numbers of HIV molecular clone. **(B)** Percentage integration activity plotted relative to WT. **(C)** Copy number of viral DNA content in the cytoplasmic PICs as measured by qPCR and calculated using a standard graph of known copy numbers of HIV molecular clone. **(D)** Percentage viral DNA content plotted relative to WT cytoplasmic PICs. **(E)** Ratio of the copy number of integrated viral DNAs to the corresponding PIC-associated viral DNAs plotted as percentage relative to WT. Data shown are representative of at least three independent experiments, with error bars representing the SEMs.

### CypA depletion restores the integration efficiency of the RK mutant virus

Interaction between HIV-1 CA and CypA is essential for optimal HIV-1 infection and any perturbation of this interaction affects reverse transcription and nuclear import^91-94, 97, 98^. Interestingly, absence of CypA in the target cell has been reported to rescue the infectivity defect of the RK mutant virus^61, 66^. The underlying mechanism behind this atypical restriction effect of CypA on the RK mutant virus infection remains unclear. Nevertheless, because our results indicated that the RK mutation impairs HIV-1 infection primarily at the integration step, we hypothesized that the rescue of the infectivity defect by the absence of CypA is a manifestation of restored integration. To test our hypothesis, we assessed reverse transcription, nuclear import and integration efficiency of the WT and the RK mutant in the parental Jurkat (Jurkat CypA+/+) and CypA knockout Jurkat (JurkatCypA-/-) cells^133^. These cells were spinoculated for 2 hours with equivalent MOI of the WT or RK mutant virus and then cultured for 24 hours, after which total DNA was isolated. The copy numbers of late RT products, 2-LTR circles, and the proviral DNA levels in the DNA samples were measured by qPCR.

The RK mutation did not alter the levels of reverse transcription (Fig. 9A and 9B) and 2- LTR circles (Fig. 9C and 9D), or the nuclear import efficiency (Fig. 9E) in the Jurkat CypA+/+ cells, thereby corroborating the data presented in Fig. 6A-E. The ∼2-fold reduction in the late RT product levels of the WT virus in the JurkatCypA-/- cells has previously been reported to be due to the lack of the positive effect of CypA on HIV-1 reverse transcription^94^. Notably, our results showing ∼2- fold reduction in the late RT product levels of the RK mutant virus in the JurkatCypA-/- cells indicate that the reverse transcription of the RK mutant virus is equally subject to the negative impact of the lack of CypA in the target cells. Interestingly, the proviral DNA copies (Fig. 9F and 9G) and the calculation of the virus integration efficiency by normalizing the proviral DNA levels to the corresponding late RT product levels and the plotting of the data as percentage relative to WT virus in parental Jurkat cells (Fig. 9H), revealed that the lack of CypA restored the proviral integration efficiency of the RK mutant virus to almost WT levels (Fig. 9F-H). These findings suggest that the rescue of the infectivity defect of the RK mutant in CypA depleted cells is primarily due to the restoration of the viral integration.

**Figure 9.**
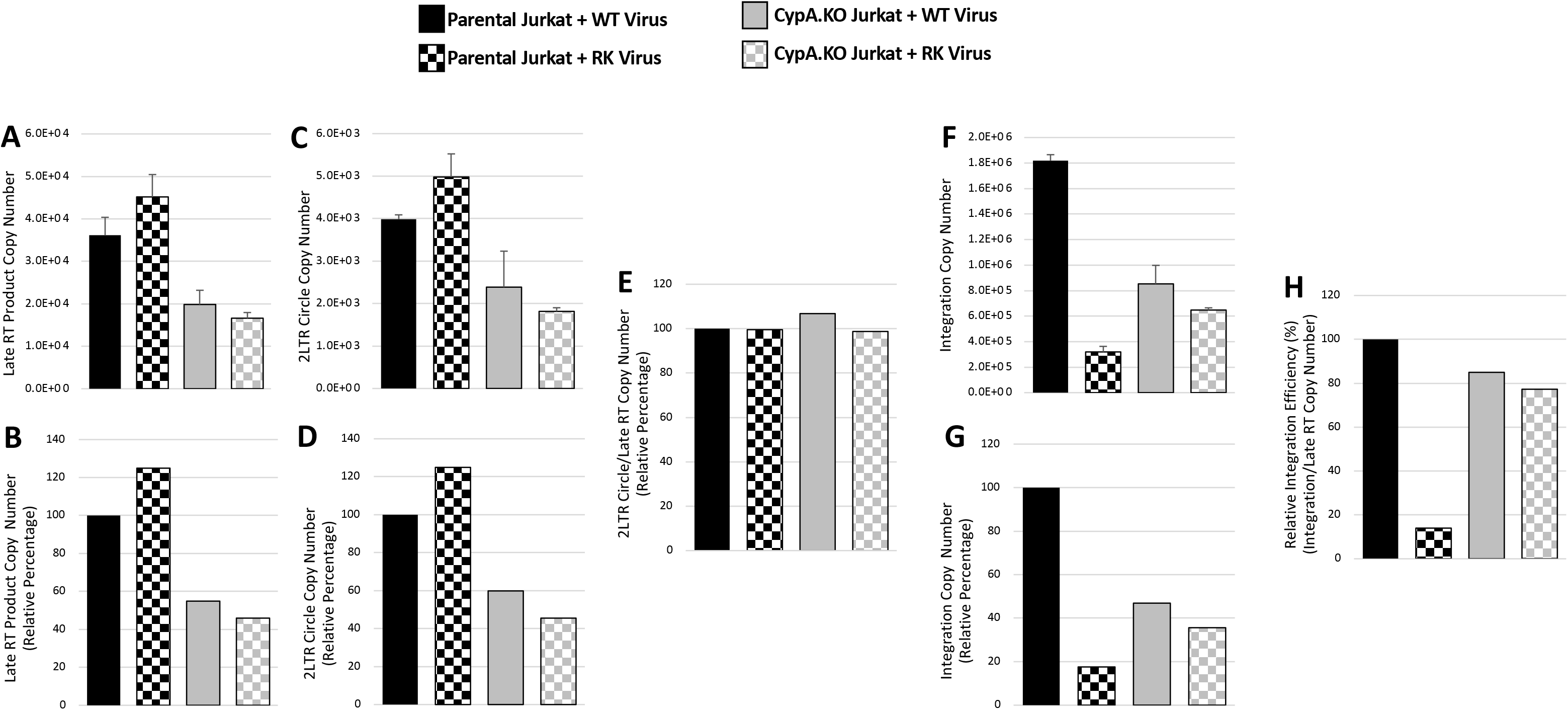
Assessment of the effect of KK10-linked CTL escape *mutation* R264K on viral DNA integration in cells lacking host cyclophilin A. Copy number of the reverse transcription (RT) products, 2-LTR circles, and the chromosome-integrated viral DNAs present in the total DNA isolated from the parental or CypA.KO Jurkat cells spinoculated with WT or the RK virus for 2 hours at 25°C and cultured for 24 hours at 37°C were determined as described in Materials and Methods. **(A)** Copy number of the late RT products in the parental or the CypA.KO cells as measured by qPCR and calculated by using a standard graph of known copy numbers of HIV-1 molecular clone. **(B)** Percentage late RT products plotted relative to WT virus in parental Jurkat cells. **(C)** Copy number of the 2-LTR circles as measured by qPCR and calculated by using a standard graph of known copy numbers of plasmid containing HIV-1 2-LTR. **(D)** Percentage 2- LTR circles plotted relative to WT virus in parental Jurkat cells. **(E)** Calculated ratio of the copy number of 2-LTR circles to the respective late RT products plotted as percentage relative to WT virus in parental Jurkat cells. **(F)** Copy number of chromosome-integrated viral DNAs as measured by *Alu*-Gag nested PCR and calculated by using a standard graph of known copy numbers of HIV-1 molecular clone. **(G)** Percentage copy number of chromosome-integrated viral DNA plotted relative to WT virus in parental Jurkat cells. **(H)** Ratio of the copy number of chromosome-integrated viral DNAs to the respective late RT products was calculated and the data plotted as percentage relative to WT virus in parental Jurkat cells. Data shown are representative of three independent experiments, with error bars representing the SEMs.

### RK mutant viral DNA ends are appended with aberrant viral sequences

Our data showing that the RK mutation significantly inhibits viral DNA integration without affecting reverse transcription levels, nuclear import efficiency, and PIC function early during infection was intriguing. We hypothesized that the reduction in viral DNA integration into nuclear chromatin becomes evident only after the PIC enters the nucleus. Therefore, we tested whether the RK mutation compromises the integrity of the viral DNA ends after the nuclear entry step of infection. The 2-LTR circles, which are generated when the open ends of the unintegrated linear viral DNAs are joined by host cell machinery^119, 120^ represent an elegant experimental tool to probe the ends of the unintegrated viral DNAs^134-136^ and thereby assess their competency for integration. Thus, we examined the nucleotide sequences of the junctions of the 2-LTR circles in cells inoculated with the WT or the RK mutant virus. The junctions of the 2-LTR circles present in the total DNA samples from duplicate infections of each virus were first amplified by nested PCR and then cloned into a plasmid vector. For each virus type, at least 100 clones were derived from two independent PCR, and the sequences of the cloned 2-LTR junctions were determined by Sanger sequencing and analyzed by multiple alignment.

Results from this analysis revealed two interesting patterns (Table 1, Fig. S4A and S4B). First, compared to the WT virus, a higher proportion of the atypically appended aberrant viral sequences at the RK mutant viral DNA ends were extensively longer. Two, a significantly higher proportion of these aberrant sequences mapped to the viral poly-purine tract (PPT) and/or the tRNA/primer-binding site (PBS) sequences. It is possible that this atypical retention of the PPT (plus-strand primer) and tRNA (minus-strand primer) sequences at the RK mutant viral DNA ends is a contributory factor in its impaired integration. These results raised the possibility that the presence of the compensatory CA mutation SA may preclude the atypical retention of such aberrant PPT and/or PBS sequences at the ends of the RKLMSA viral DNAs. Remarkably, compared to the RK mutant viral DNAs, the proportion of the RKLMSA mutant viral DNAs with aberrant PPT and/or PBS sequences were significantly low and were even lesser than the WT sequences (Table 1, Fig. S4C). Together, these results suggest that the retention of the PPT and PBS sequences at the ends of the RK mutant viral DNAs may be contributing to their impaired integration and may also be indicative of an underlying qualitative glitch in reverse transcription (Figure 10).

**Table 1.** Assessment of the effect of KK10-linked CTL escape mutations on the integrity of the termini of the reverse transcription products (viral DNAs). Junctions of the 2-LTR circles present in the total DNA isolated from Jurkat cells spinoculated with the WT or mutant viruses for 2 hours at 25°C and cultured for 24 hours at 37°C were amplified by a nested PCR strategy, cloned into a plasmid vector, and the DNA sequences determined by Sanger sequencing were aligned and analyzed as described in Materials and Methods. The total number of 2-LTR junction sequences analyzed for the WT and the mutant viruses is indicated at the top of the table. The total number of each type of 2-LTR junction sequence identified in the analyses is shown for the WT, RK, and RKLMSA virus-inoculated samples. Shown at the bottom of the table are the total number of 2-LTR junctions containing aberrant insertion sequences and their percentage relative to the total sequences analyzed, and the total number of insertion sequences mapping to the PPT or PBS and their percentage relative to the total insertion sequences. See also Supplementary Figure S4.

|  |  | WT | RK | RKLMSA |
| --- | --- | --- | --- | --- |
| **Total Number of Sequences Analyzed** |  | 104 | 106 | 166 |
| **2-LTR junctions without aberrant nucleotide insertions** | Complete 3'-end processing of both LTR ends, with intact flanking regions | 0 | 0 | 0 |
|  | Complete 3'-end processing of both LTR ends, accompanied by deletions in flanking regions | 6 | 14 | 15 |
|  | Complete 3'-end processing of one LTR end (5'U3 or 3'U5) | 1 | 7 | 14 |
|  | No 3'-end processing of both LTR ends | 15 | 7 | 13 |
|  | Partial 3'-end processing of one LTR end | 20 | 12 | 10 |
| **2-LTR junctions with aberrant nucleotide insertions** | Complete 3'-end processing of both LTR ends, with intact flanking regions | 0 | 1 | 1 |
|  | Complete 3'-end processing of both LTR ends, accompanied by deletions in flanking regions | 2 | 1 | 8 |
|  | Complete 3'-end processing of one LTR end (5'U3 or 3'U5) | 21 | 32 | 34 |
|  | No 3'-end processing of both LTR ends | 35 | 26 | 59 |
|  | Partial 3'-end processing of one LTR end | 5 | 6 | 22 |
| **Source of aberrant insertion sequences** | Number of clones with insertions at the junction (% of total sequences) | 63 (60.6%) | 66 (62.3%) | 119 (71.7%) |
|  | Insertions mapping to PPT sequences (% of total insertion sequences) | 4 (6.3%) | 16 (24.2%) | 7 (5.9%) |
|  | Insertions mapping to PBS sequences (% of total insertion sequences) | 8 (12.7%) | 25 (37.9%) | 9 (7.6%) |

**Figure 10.**
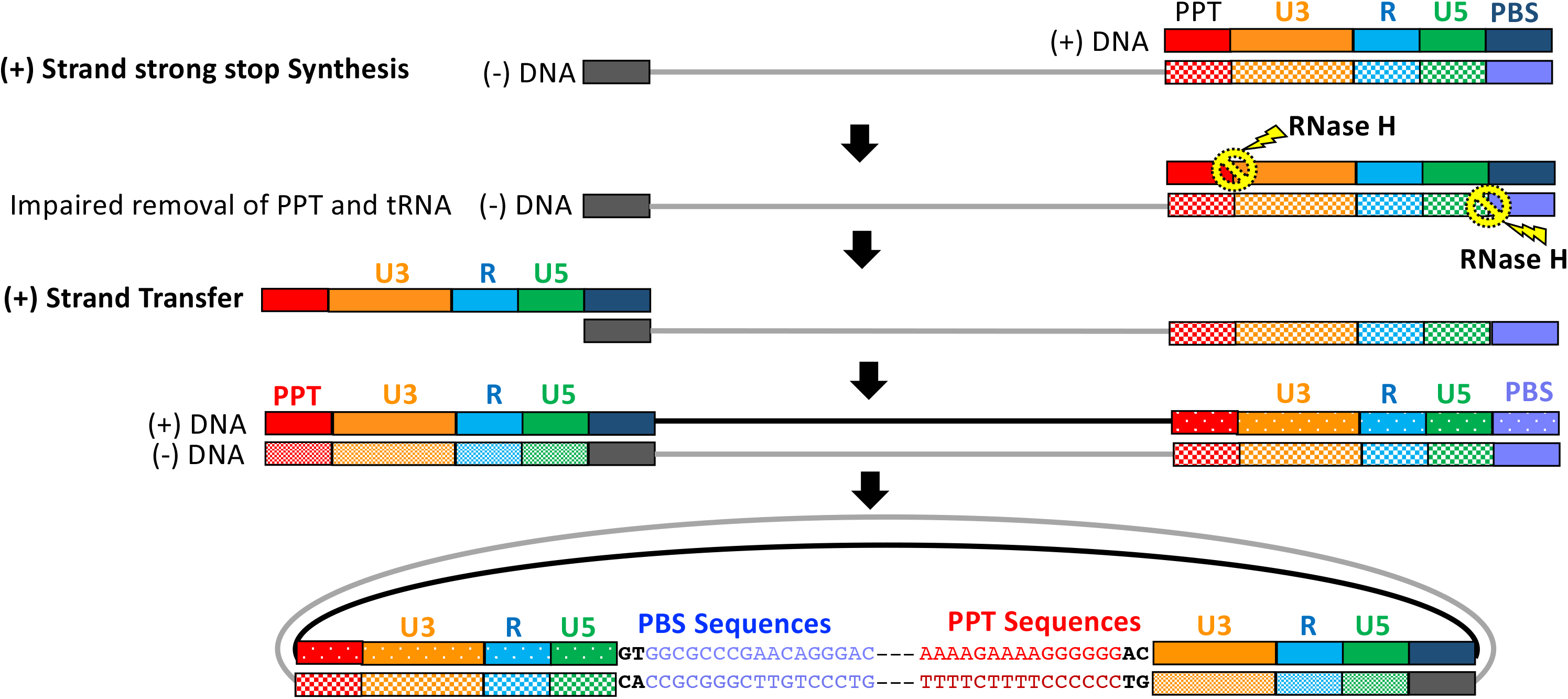
Model for the presence of PPT and PBS sequences at HIV-1 DNA ends. Shown schematically is HIV-1 reverse transcription after the completion of the plus-strand strong stop DNA synthesis. In the case of wild type HIV-1, the RNase H removes the PPT/plus-strand primer (at the junction between PPT and plus-strand strong stop DNA) and the tRNA/minus-strand primer (one base from the junction between tRNA and minus-strand DNA). These RNase H-mediated removal of the PPT and tRNA primers, though not necessary for strand transfer, are essential for the formation of proper viral DNA ends that, after the synthesis of the complete double-stranded viral DNA, can function as competent substrates for the integration reaction. However, in the case of the RK mutant HIV-1, increased failure of RNase H to properly remove the PPT and tRNA primers can lead to the increased retention of these primer sequences at the respective viral DNA ends. These viral DNA ends are unsuitable substrates for the viral integrase-mediated integration reaction in the nucleus and therefore are ligated by host enzymes to form 2-LTR circles.

## DISCUSSION

The human immune response against the founder HIV-1 is executed by the CTLs in an HLA genotype-dependent manner. The CTL response against the HIV-1 KK10 epitope has been linked to robust virus control and delayed disease progression in certain HLA-B27-positive infected individuals. Selection of the CA mutation RK, associated with viral escape, disrupts epitope binding to HLA-B27 but also causes a severe infectivity defect. The compensatory CA mutation SA relieves the RK-associated virus infectivity defect and triggers loss of viremia control and progression to AIDS. In this study, we have identified impaired viral DNA integration as the principal molecular mechanism underlying the infectivity defect of the RK mutant virus. Our data also establishes that the RK mutation has limited impact on the efficiency of reverse transcription and nuclear import steps of virus replication and the SA compensatory mutation largely rescues the integration defect of the CTL-escape RK mutant virus.

Specific mutations in the genetically fragile CA compromise capsid integrity and disrupt early events of HIV-1 replication. While Schneidewind et al.^61^ reported that the RK mutation had no significant effect on capsid stability, work by Schommers et al.^67^ revealed a modest increase in the stability of the RK mutant capsid. It is noteworthy that the RK mutation in the CA does not appear to overlap with the critical structural elements of the capsid. Accordingly, our molecular modeling and MD simulations demonstrated that the RK mutation neither altered the integrity of the hexameric, pentameric or the tubular structure of the capsid. Further, the RK mutation minimally affected interaction with CypA- a critical host factor known to modulate the capsid stability. Our Y2H assays corroborated these findings revealing that the binary interaction between the RK CA or the RK CA and CypA are largely preserved.

Our results confirmed that the RK mutation—independently or in combination with the LM mutation—causes a drastic loss of infectivity but could be rescued by the compensatory SA mutation. Interestingly, our infection assays carried out in an epithelial cell line (TZM-bl) and in a T-cell line (Jurkat) with native envelope-containing viruses inform that the consequence of the RK, RKLM, and RKLMSA mutation on infectivity are unlikely to be dependent on cell types. Further, the infectivity defect of these variants irrespective of whether the virus contains the native HIV-1 envelope or the VSVg-pseudotyped envelope^51, 67^ suggests that the mode of viral entry, i.e., fusion vs endocytosis, does not influence how these CA mutations impair infectivity. Our results of the Gag processing and virus release efficiency also preclude effect of these mutations on late stages of infection. Therefore, the RK-associated infectivity defect most likely arises from impairment of one or more of the early post-entry steps of infection.

Our findings that the RK and RKLM mutations are unlikely to perturb the integrity of the capsid and its interaction with CypA suggested that these CA mutations may not affect the reverse transcription efficiency. However, Schneidewind et al^61^ reported that the RK mutation, despite having no effect on the capsid stability, caused a severe reduction in HIV-1 reverse transcription levels. In contrast, our quantification of reverse transcription levels did not reveal any significant difference between the WT and the RK or RKLM mutant viruses. While Schneidewind et al.^61^ investigated reverse transcription using VSVg-pseudotyped virus that enters target cells by endocytosis, we measured the reverse transcription levels using native envelope-containing RK mutant virus that enters target cells by CD4-receptor mediated membrane fusion. This raises the possibility of a pseudotyping-associated effect on the RK mutant virus reverse transcription, since VSVg pseudotyping has been reported to alter the phenotype of certain CA mutants^137, 138^.

The lack of a significant effect of the RK mutation on HIV-1 reverse transcription levels in our study implies that this CA mutation disrupts one or more of the downstream steps in HIV-1 infection. Our quantification of the 2-LTR circles, widely considered a reliable surrogate marker for HIV-1 nuclear import, showed no significant differences in the levels of nuclear import of the WT or RK or RKLM mutant virus. It is increasingly becoming clear that the interaction between the HIV-1 CA and the host CPSF6 is essential for optimal nuclear import of the virus. Interestingly, results from our molecular modeling, MD simulations, and yeast two-hybrid assays showing no significant effect of the RK or RKLM mutations on the interaction between CA and CPSF6 further supports our 2-LTR circle data and the improbability of a RK mutation-induced defect in viral nuclear import. Accordingly, measurement of the proviral copy numbers revealed that the CTL escape CA mutations RK and RKLM severely impaired the ability of the viral DNA to integrate into the host chromatin. Importantly, the compensatory SA mutation was able to significantly rescue the viral DNA integration defect exhibited by the RK and RKLM mutant viruses. Collectively, these results provide strong evidence that the RK mutation-associated reduction in infectivity is not due to alterations in reverse transcription and nuclear import but exclusively a consequence of impaired viral DNA integration.

In infected cells, integration of the viral DNA into the host chromosome is carried out by the HIV-1 PIC. Therefore, we hypothesized that a diminished integration activity of the RK PICs could be the underlying cause of impaired viral DNA integration of the mutant virus. Our results showed that the viral DNA levels in the isolated WT and RK mutant cytoplasmic PICs were comparable, thereby further corroborating that the RK mutation has no significant effect on viral reverse transcription levels. Unexpectedly, the integration activity of the WT and RK mutant cytoplasmic PICs did not vary significantly, in striking contrast to a significant reduction in the number of RK mutant proviral DNAs. There are key differences in our assays measuring *in vitro* integration and the proviral integration. These include contrasting nature of the target DNAs (naked DNA vs chromosomal DNA), distinct composition of the cytoplasmic PICs and nuclear PICs (e.g., reported reduction in PIC-associated integrase and CA levels upon nuclear entry), and spatial and temporal constraints in the involvement of cellular cofactors (e.g., nuclear CPSF6 and LEDGF). Although these differences may play a role, it should be noted that our measurements of reverse transcription are quantitative but not qualitative. Furthermore, emerging evidence suggest that HIV-1 reverse transcription may not be completed until the PIC enters the nucleus. Because proviral integration is dependent on precise viral DNA ends, we investigated whether the quality of the viral DNAs in the nucleus are incompetent for integration.

The sequence information of the junctions of 2-LTR circles, which are typically generated in the nucleus of HIV-1-infected cells by the end-to-end ligation of unintegrated viral DNA molecules, can be used to assess the integration competency. A key step leading up to HIV-1 integration is the 3’-end processing of viral DNAs containing precisely-defined termini. The generation of viral DNA with canonical termini depends on completion of strand transfer reactions and the proper removal of the tRNA and the PPT primers used for the minus strand and plus strand DNA synthesis, respectively. Any disruption of the above events results in viral DNA termini that are incompetent for integration and, consequently, are circularized by host machinery into 2- LTR circles. Accordingly, the 2-LTR circle junction sequences are typified by the presence of unprocessed viral DNA ends, deletions within the 3’U5 or 5’U3 region, and insertion (i.e., retention) of aberrant viral DNA mapping mostly to PPT and PBS sequences. Indeed, we observed all these modifications in our comparative analysis of the cloned junction sequences of 2-LTR circles from cells inoculated with the WT or RK mutant virus. However, compared to the WT, the insertions in RK 2-LTR circle junctions presented two distinctive patterns. One, a higher proportion of these insertions were significantly longer. Two, a significantly higher proportion of these insertions mapped to the viral PPT and/or the PBS sequences. This atypical and enhanced retention of the PPT and PBS sequences at the RK mutant viral DNA termini could be a major factor contributing to its impaired integration. Remarkably, compared to the RK mutant viral DNAs, the proportion of the RKLMSA mutant viral DNAs atypically appended with aberrant PPT and/or PBS sequences were significantly low and were even lesser when compared to the WT sequences. The atypical retention of the PPT and PBS sequences at viral DNA termini has previously been proposed to arise from non-optimal RNase H activity of the viral reverse transcriptase or from autointegration. It is unclear how the RK mutation that is positioned in the CA and did not cause any significant alterations in the reverse transcription levels can selectively alter the RNase H activity of the reverse transcriptase. Further, autointegration is unlikely to be the source of these viral DNA insertions for three reasons. One, primers used in the PCR-based method for obtaining the 2-LTR circle junction sequences are complementary only to 2-LTR circle- specific sequences. Two, our use of 24-hpi sample as the source of template for PCR excludes the detection of autointegrants because they accumulate at early time points (∼10 hpi)^139^ and are rapidly depleted by 24 hpi. Three, requirement of viral DNA 3’-end processing for the integrase-mediated autointegration is incompatible with our sequencing data showing that majority of the viral DNA termini are unprocessed or partially processed. Nevertheless, these results suggest that the atypical retention of the PPT and PBS sequences at the ends of the RK mutant viral DNAs may significantly contribute to their drastically impaired integration into chromosomal DNA and suggest an underlying qualitative glitch in reverse transcription.

The rescue of the infectivity defect of the RK mutant virus in CypA-depleted cells and in cells treated with cyclosporin A indicates that an interaction between the RK mutant CA and CypA is a prerequisite for the manifestation of the virus infectivity defect^61, 67^. Accordingly, results from our virus infection assays carried out in CypA-depleted cells indicated that the infectivity of the RK mutant virus was comparable to that of the WT virus. Is CypA playing a causative role in the infectivity defect incurred by the RK mutant virus? Interestingly, supportive evidence comes from our data comparing the viral DNA integration levels of the RK mutant virus with that of the WT virus in the CypA knockout Jurkat cells. In line with the infectivity assays, the viral DNA integration level of the RK mutant virus was comparable to that of the WT virus in the CypA knockout Jurkat cells. Especially striking is the several fold increase in the integration levels of the RK mutant virus in the CypA knockout Jurkat cells relative to the control Jurkat cells. This indicates that CypA expression and its binding to the mutant CA is a prerequisite for the significant reduction in viral DNA integration by the RK mutant virus. Does this also imply that the reduction in virus infectivity by RK mutation is exclusively a consequence of the CypA-associated impaired viral DNA integration? In line with the findings from the control Jurkat cells, the reverse transcription and nuclear import levels of the RK mutant virus was comparable to that of the WT virus in the CypA knockout Jurkat cells. This indicates that the CypA knockout-mediated restoration of the RK mutant virus infectivity is primarily an outcome of the rescue of the RK mutation-induced integration defect. It should be noted that the prior studies did not assess the effect of these CA mutations on HIV-1 integration^61, 67^ and, instead, the log-fold reduction in infectivity of the RK and RKLM mutant viruses in single-cycle infectivity assays were attributed to the reduction in the levels of reverse transcription^61^.

This work also raises several important questions, including: (1) How does the RK mutation lead to the generation of imperfect viral DNA ends? (2) Why is CypA expression a requirement for the RK mutation-associated integration defect? and (3) How does the compensatory CA mutation SA restore the RK mutant virus infectivity without disrupting the interaction between the mutant CA and host CypA? The RK mutation appears to neither disrupt the interaction between CA and host CypA nor alter the quantitative aspects of reverse transcription. However, it is possible that the interaction between the RK mutant CA and the host CypA leads to an atypical and prolonged association between these proteins. This, in turn, by altering qualitative aspects of reverse transcription (e.g., altered RNase H activity) may lead to the generation of an increased proportion of viral DNA containing integration-incompetent termini. Conversely, the compensatory CA mutation SA or absence of CypA may preclude such an atypical and prolonged association between the RK mutant CA and CypA. However, the ability of the compensatory SA mutation to restore the RK mutant virus integration without imposing any loss of interaction between the mutant CA and host CypA, suggest the possible involvement of additional cellular co-factor(s). Accordingly, CPSF6 expression was recently reported as a requirement for the manifestation of the RK mutation-associated virus infectivity defect^66^. Although there is no evidence linking the interaction between CA and CPSF6 to reverse transcription, a recent study reported that CypA binding to HIV-1 capsid prevents oligomerized cytoplasmic CPSF6 from prematurely binding and disrupting the capsid^140, 141^. An alternative possibility, from our in vitro integration assays showing that the cytoplasmic PICs are competent for viral DNA integration, is that the RK mutation-associated aberrant viral DNA termini are generated only in the nucleus. For instance, an atypical association between RK mutant CA and CypA may delay viral integration during which the viral DNA termini may be subjected to host DNA polymerase-mediated copying and insertion reactions, thereby resulting in the generation of aberrant viral DNA termini in the nucleus.

It is important to note that ART only keeps HIV-1 replication in check and does not cure or eradicate the lifelong infection. One approach called “functional cure” entails controlling the virus to undetectable levels without ART and avert disease progression to AIDS^142, 143^. Remarkably, the elite controllers possess all these characteristics^22, 144, 145^ and are considered a suitable model for achieving functional cure^24, 146-148^. The elite controller phenotype has been linked to robust killing of the HIV-1-infected cells by CTLs^149^ and has been shown to be enriched in individuals with certain HLA genotypes, including HLA-B27. Our work, showing impaired HIV- 1 integration as the driver of the fitness cost of the clinically relevant CTL escape RK mutation (associated with HLA-B27), provides a novel mechanistic basis to advance CTL-based HIV cure strategies^37, 52, 150-153^ and to rationally design superior epitopes for T cell-based HIV vaccines.

## MATERIALS AND METHODS

### Cell culture

The human embryonic kidney cell line HEK293T and the T-cell line SupT1 were obtained from the American Type Culture Collection (Manassas, VA, USA). The following cell lines were obtained through the NIH AIDS Reagent Program, Division of AIDS, NIAID, NIH (USA): Jurkat (E6-1) and Jurkat CypA deficient (Jurkat CypA-/-) T-cell lines from Douglas Braaten and Jeremy Luban, and the TZM-bl reporter cell line from John C. Kappes, Xiaoyun Wu, and Tranzyme, Inc. The HEK293T and TZM-bl cells were cultured in Dulbecco modified Eagle medium (DMEM; Gibco/Thermo Scientific, USA) supplemented with 10% heat-inactivated fetal bovine serum (FBS; Gibco/Thermo Scientific, USA), 2 mM glutamine, 100 U/mL penicillin, and 100 μg/mL streptomycin. The SupT1, Jurkat, Jukat CypA-/- cell lines were cultured in Roswell Park Memorial Institute (RPMI) 1640 medium supplemented with 10% heat-inactivated FBS, 2 mM glutamine, 100 U/mL penicillin, and 100 μg/mL streptomycin. Cells were cultured at 37°C with 5% CO2.

### Proviral plasmids and cloning

The viruses used in this study were generated from the full-length wildtype HIV-1 molecular clone pNL43 and its mutant derivatives. Unless indicated otherwise, all PCRs were performed using the high-fidelity Phusion DNA polymerase (NEB, USA) and custom-made primers. Agarose gel-resolved DNAs were extracted using the Zymoclean Gel DNA Recovery Kit (Zymo Research, USA); plasmid DNA and PCR amplicons were digested with commercial restriction enzymes (NEB, USA); DNAs were ligated using T4 DNA ligase (NEB, USA); competent DH5α (Invitrogen, USA) or NEB-Stbl (NEB, USA) bacterial cells were used for bacterial transformation; plasmid DNAs were isolated using the Zyppy plasmid mini prep kit or ZymoPURE II plasmid maxiprep kit (Zymo Research, USA); recombinant plasmids were verified by restriction enzyme digestion and Sanger DNA sequencing; and site-specific mutations were introduced into DNA using the Q5 Site-Directed Mutagenesis Kit (NEB, USA) and custom-made primers. Site-specific mutations in the capsid (CA) coding region of pNL43 were first introduced into a 1310-bp DNA fragment—spanning the matrix (MA), capsid (CA), and part of the nucleocapsid (NC) coding regions and flanked by the BSSHII and ApaI restriction sites— of pNL43 cloned into the pMiniT 2.0 vector (NEB, USA). This pMiniT2-Gag/BssHII-ApaI intermediary plasmid was constructed by cloning a PCR amplicon obtained by using custom-made forward primer harboring BssHII site (5’- TATAGCGCGCACGGCAAGAG-3’) and reverse primer harboring ApaI site (5’- TATAGGGCCCTGCAATTTTTGGCTATG-3’) of the corresponding DNA sequence from pNL43 into the linearized pMiniT 2.0 vector (NEB, USA) as per manufacturer-recommended protocol. The R264K/G791A mutation was introduced using a forward primer containing the desired mutation (G>A: 5’-ATCTATAAAAAATGGATAATCCTG-3’) and a reverse primer (5’-TTCTCCTACTGGGATAGG-3’) to yield the pMiniT2-Gag-RK plasmid. The L268M/C802A mutation was introduced using a forward primer containing the desired mutation (C>A: 5’- ATGGATAATCATGGGATTAAATAAAATAG-3’) and a reverse primer (5’- CTTTTATAGATTTCTCCTACTG-3’) to derive the pMiniT2-Gag-LM plasmid. The R264K/G791A mutation was introduced into the pMiniT2-Gag-LM plasmid using a forward primer containing the desired mutations (G>A: 5’-AGGAGAAATCTATAAAAAATGGATAATCATGG-3’) and a reverse primer (5’- ACTGGGATAGGTGGATTATGTG-3’) to yield the pMiniT2-Gag-RKLM plasmid. The S173A/T517G mutation was introduced into the pMiniT2-Gag-RKLM plasmid using a forward primer containing the desired mutations (G>A: 5’-ACCCATGTTTGCAGCATTATC-3’) and a reverse primer (5’- ATTACTTCTGGGCTGAAAG-3’) to yield the pMiniT2-Gag-RKLMSA plasmid. The BssHII-ApaI DNA fragments from the pMiniT2-Gag-RK, pMiniT2-Gag-LM, pMiniT2-Gag- RKLM, and pMiniT2-Gag-RKLMSA plasmids were released by digestion with the BssHII and ApaI enzymes, and the gel-purified DNA fragments were then individually ligated to the BssHII+ApaI- cut pNL43 plasmid to yield pNL43-RK, pNL43-LM, pNL43-RKLM, and pNL43-RKLMSA plasmids, respectively.

### GAL4-based yeast two-hybrid assay

The GAL4-based yeast two-hybrid (Y2H) system parental plasmids, pGBKT7 and pGADT7 (Clontech, USA), were used to generate the Y2H plasmids encoding the viral or host proteins. The DNA sequence encoding the CA ORF was amplified from the pNL43 plasmid by PCR using a forward primer harboring EcoR1 site (5’- TAAGCAGAATTCCCTATAGTGCAGAACCTCCAGG-3’) and a reverse primer harboring Sal1 site (5’-TCATTAGTCGA*CTATCA*CAAAACTCTTGCTTTATGG-3’) and GoTaq DNA polymerase (Promega, USA). The PCR amplicon was gel-purified using the Qiaquick gel purification kit (Qiagen, USA), digested with EcoR1 and Sal1 enzymes, and then ligated to the EcoR1+Sal1-cut pGBKT7 plasmid or EcoR1+Xho1-cut pGADT7 plasmid to yield the pGBKT7-CA and pGADT7-CA plasmids, respectively. The DNA sequence encoding the mCherry ORF was amplified from mCherry-H2A-10 (Addgene plasmid# 55054) by PCR using a forward primer harboring EcoR1 site (5’-TGACAGAATTCATGGTGAGCAAGG-3’) and a reverse primer harboring the Xho1 site (5’-TGACACTCGAGTTACTTGTACAGC-3’). The gel-purified PCR amplicon was digested with EcoR1 and Xho1 enzymes and then ligated to the EcoR1+Sal1-cut pGBKT7 plasmid or EcoR1+Xho1-cut pGADT7 plasmid to yield the pGBKT7-mCherry and pGADT7-mCherry, respectively. The DNA sequence encoding the CypA ORF was released from the pET21-CypA plasmid by digestion with Nde1 and Xho1 enzymes, and the gel-purified Nde1+Xho1-cut CypA ORF was ligated to Nde1+Xho1-cut pGADT7 plasmid to yield the pGADT7-CypA. Site-specific mutations in the DNA sequence encoding the CA ORF in the pGBKT7-CA and pGADT7-CA plasmids were introduced using custom-made primers. The R264K/G791A mutation was introduced using a forward primer containing the desired mutation (G>A: 5’- ATCTATAAAAAATGGATAATCCTG-3’) and a reverse primer (5’-TTCTCCTACTGGGATAGG-3’) to yield the pGBKT7-CA-RK and pGADT7-CA-RK plasmids, respectively. The DNA sequence encoding the CPSF6 ORF was amplified from the pTarget-CPSF6 by PCR using a forward primer harboring EcoR1 site (5’- TATAGAATTCATGGCGGACGGCGTGG-3’) and a reverse primer harboring the Xho1 site (5’-TATACTCGAGCTAACGATGACGATATTCGCGCTC-3’). The gel-purified PCR amplicon was digested with EcoR1 and Xho1 enzymes and then ligated to the EcoR1+Xho1-cut pGADT7 plasmid.

Evaluation of protein interactions by the GAL4-based Y2H system was performed by following the manufacturer’s recommendations for the Matchmaker GAL4 Two-hybrid System 3 (Clontech, USA). Competent AH109 yeast cells (Clontech, USA) were prepared and transformed with the Y2H plasmid constructs by using the Frozen-EZ Yeast Transformation II kit (Zymo Research, USA) and following the manufacturer-supplied protocol and recommendations. Briefly, competent AH109 cells were co-transformed with appropriate combinations of bait and prey Y2H plasmids and selected for growth on minimal synthetic dropout (SD) agar plates lacking tryptophan (-T) and leucine (-L) [SD/-T/-L] at 30°C. The ability of the bait and prey proteins to interact was assayed by testing the yeast transformant’s ability to grow on SD agar plates lacking tryptophan, leucine, and histidine (SD/-T/-L/-H) at 30°C, which is indicative of a positive protein interaction-driven expression of the reporter gene *HIS3*. Positive protein-protein interactions were subjected to another round of stringent selection by testing the yeast transformant’s ability to express *α*-galactosidase from the *MEL1* reporter gene (blue/white screening) when selected on SD/-T/-L/-H agar plates supplemented with X-*α*-Gal (Sigma, USA) at 30°C.

### Molecular Modelling Studies

Molecular modelling of the CA hexamer structure: The initial coordinates of the HIV-1 CA hexamer were generated by applying a six-fold symmetry operation onto a native full-length HIV- 1 capsid protein (PDB accession code 4XFX). The two loops between residues 5 to 9 and residues 222 to 231, missing in the original structure, were built by Modeler ^154^. Three models of CA hexamer mutations, R132K (RK), R132KL136M (RKLM), and R132KL136MS41A (RKLMSA), were generated in VMD^155^ Mutator Plugin from the 4XFX hexamer model. After placed an IP6 molecule^106^ close to the Arg18 ring, sodium and chloride ions were introduced around these models, based on the local electrostatic potential, using CIONIZE plugin in VMD. Subsequently, all models were then solvated with CHARMM TIP3P water model^156^ and the total NaCl concentrations were set to 150 mM, resulting charge-neutral systems of about 65 K atoms.

Molecular modelling of the CA pentamer structures: Details about the molecular modeling of two CA pentamer structures were described in Xu, et al. (2020)^107^. Briefly, the coordinates of the disulfide stabilized pentamer were from crystal structure 3P05, after mutating the cysteine residues back to the HIV-1NL4-3 wildtype sequence. The coordinates of the missing residues in the PDB model were built by Modeler^154^. Using the mutated 3P05 pentamer structure for the initial coordinates, the 5MCY pentamer model was generated by running molecular dynamics flexible fitting (MDFF) of the initial coordinates into 5MCY cryo-EM density (EMBD: EMD-3466).

Simulation protocol: The solvated systems were then subjected to minimization in two stages, both using the conjugated gradient algorithm^157^ with line search^158^. Each stage consisted of 10,000 steps of energy minimization. During the first stage, only water molecules and ions were free to move, while the protein and IP6 molecule were fixed. In the second stage, the backbone atoms of the CA protein were applied with a harmonic restraint with a force constant of 10.0 kcal mol^-1^ Å^-2^. Convergence of the minimization procedure was confirmed once the variance of the gradient was below 0.1 kcal mol^-1^ Å^-1^. Following minimization, the systems were tempered from 50 K to 310 K in increments of 20 K over 1 ns. Subsequently, the systems were equilibrated at 310 K for 100,000 steps, while the protein backbone atoms were restrained. Then positional restraints were gradually released at a rate of 1.0 Kcal mol^-1^ Å^-2^ per 400 ps from 10.0 Kcal mol^-1^ Å^-2^ to 0.0 Kcal mol^-1^ Å^-2^.

All MD simulations in present study were performed with NAMD2.13^159^ using CHARMM force fields^160, 161^. In present study, an internal time step of 2 fs was employed in the multi-step vRESPA integrator as implemented in NAMD, bonded interactions were evaluated every 2 fs. Temperature was held constant at 310 K using a Langevin thermostat with a coupling constant of 0.1 ps^-1^. Pressure was controlled at 1 bar using a Nose-Hoover Langevin piston barostat with period and decay of 40 ps and 10 ps, respectively. The Shake algorithm was employed to constraint vibrations of all hydrogen atoms. Long range electrostatics was calculated using the particle-mesh-Ewald summation with a grid size of 1 Å and a cutoff for short-range electrostatics interactions of 12 Å. MD result analysis: The RMSD, RMSF, contact and ion occupancy analyses were performed in VMD^155^. The APBS software^162^ was employed to compute the electrostatic potential surfaces. The sodium and chloride ions were added to all systems according to the local coulombic potential by CIONIZE in VMD^155^.

### Cell- and virus-associated viral protein expression in cells transfected with proviral plasmids

HEK293T cells, seeded at 6 × 10^5^ cells per well in a 6-well plate and cultured overnight, were transfected with 2 μg of WT and mutant proviral plasmid DNAs using PEI. After 24 h or 48 h post transfection, the culture media containing the released virus particles (virus fraction) was collected, and the cells were lysed in TX-100 lysis buffer (300mM NaCl; 50mM Tris-HCl, pH 7.5; 0.5% Triton X-100; β-mercaptoethanol, Sigma protease inhibitor cocktail) for 10 min on ice. The virus fraction was centrifuged at 500x *g* for 3 min to remove cells/debris, passed through 0.45-μm filter, and centrifuged in a SW 32.1 Ti rotor (Beckman Coulter) at 32,000 rpm for 45 min at 4°C. The supernatant was carefully removed, and the virus pellet was lysed in 0.1 mL of TX-100 lysis buffer for 10 min on ice. The total protein concentrations of the cell lysates were quantified by BCA protein assay (Pierce, USA). Equal amounts of the cell lysate protein and equal volume of the virus pellet protein were resolved by SDS-PAGE (NuPAGE 4-12% Bis-Tris gel; Invitrogen/Thermo Scientific) and transferred onto nitrocellulose membrane by semidry transfer. Membranes blocked in non-fat dry milk were first probed with mouse anti-CA monoclonal antibody (1:1000 dilution; 183-H12-5C; NIH AIDS Reagent Program, USA), followed by secondary HRP-conjugated goat anti-mouse IgG(H+L) (1:10,000 dilution; Bio-Rad, USA), and then developed using the enhanced chemiluminescence procedure (Pierce, USA). Virus release efficiency is calculated as the amount of virus-associated CA divided by the total Gag (virus-associated CA + cell-associated Pr55Gag + cell-associated p41Gag + cell-associated CA).

### Preparation of virus stocks and determination of titer

Virus stocks were generated by PEI-mediated transient transfection of HEK293T cells with WT or mutant HIV-1 proviral plasmid constructs. Briefly, 2 × 10^6^ HEK293T cells, seeded per 10- cm culture dish and cultured overnight, were transfected with 10 or 20 μg of plasmid DNA. After 12-16 h, the cell culture media was removed, the cells washed once with 1X phosphate-buffered saline (PBS), 6 mL of new growth medium was added to the cells, and the cells were cultured for additional 48 h. Then, the culture media containing the released virus particles was harvested, centrifuged at low-speed to remove the cell debris, passaged through 0.45-μm-pore-size syringe filters, and treated with DNase I (Calbiochem/MilliporeSigma, USA; 20 μg/mL of supernatant) in the presence of 10 mM magnesium chloride for 1 h at 37°C to eliminate any carryover plasmid DNA from transfection.

The titer of the virus preparations was determined by using the qPCR lentivirus titration kit (Applied Biological Materials, USA) as per the manufacturer-recommended protocol. Briefly, the viral RNA from the virus sample was extracted by mixing 2 μL of virus preparation with 18 μL of virus lysis buffer and incubating at room temperature for 3 min. The resulting virus lysate, and two lentiviral standards- STD1 and STD2, were used as templates, along with the reagent-mix, lentivirus 2X qPCR mastermix, and 1X iTaq Universal SYBR Green Supermix (Bio-Rad, USA), in qRT-PCRs carried out under following cycling conditions: initial incubations at 42°C for 20 min (reverse transcription) and 95°C for 10 min (enzyme activation), followed by 30 cycles of amplification and acquisition at 95°C for 15 s and 60°C for 1 min. The resulting Ct values were then used to calculate the viral titer of the virus preparations using the formula 5 x 10e7/2e3(Ctx- Ct1)/(Ct2-Ct1), where Ctx = average of 3 Ct values of the unknown sample, Ct1 = average of 3 Ct values of STD1, and Ct2 = average of 3 Ct values of STD2.

### TZM-bl-based HIV-1 infectivity assay

TZM-bl cells, seeded at 4 × 10^4^ cells per well in 24-well plate and cultured overnight, were inoculated with WT or mutant viruses (MOI of 0.05 or 1.0) in the presence of polybrene for 2 h at 37°C/5% CO2. After sequential washes with 1X PBS and cell culture medium, the cells were replenished with new complete growth medium and cultured further for 48 h. The cells in each well were washed with 1X PBS, lysed in 200 μL of 1X GLO lysis buffer (Promega, USA), and incubated with shaking at room temperature for 5 min. Luminescence activity in the resulting cell lysates was measured using the Luciferase Assay System (Promega, USA).

### Assessment of HIV-1 infection in T-cells

Jurkat cells (2.5 × 10^5^ cells per well) in 48-well plates were spinoculated with virus stocks (MOI of 1.0) at 480 × *g* for 2 h at 25°C. After the spin, the spinoculated samples were individually transferred onto 1.7 mL microcentrifuge tubes, centrifuged at 300 × *g* for 10 min at 25°C, and the supernatants were carefully removed and discarded. Each cell pellet was then thoroughly resuspended in 1 mL of RPMI complete medium, transferred onto new 48-well cell culture plate, and cultured for 24 h or 48 h at 37°C. The cells were then harvested by transferring the cell cultures to 1.7-mL microcentrifuge tubes and centrifuging at 500 × *g* for 5 min at 25°C. The cell pellets were washed once with PBS, resuspended in 300 μL of BD Cytofix/Cytoperm (BD Biosciences, USA), and incubated for 20 min at 4°C. The cells were then centrifuged at 500 × *g* for 5 min at 25°C and the supernatants were discarded. The cell pellets were washed once with 300 μL of BD Perm/Wash buffer and then resuspended in 300 μL of BD Perm/Wash buffer (BD Biosciences, USA) for storage at 4°C. The cell samples were centrifuged and the cell pellets were resuspended in 110 μL of BD Perm/Wash buffer. For staining Gag protein, 50 μL of each cell sample was mixed with 1 μL of KC57-FITC antibody that identifies the HIV-1 Gag and p24 (Beckman Coulter Life Sciences, USA) and incubated at 4°C for 20 min. The cell samples were centrifuged and the cell pellets were washed twice with 300 μL of BD Perm/Wash buffer. The cells were then resuspended in 100 μL of BD Perm/Wash buffer and the percentage of FITC-labelled cells, as a marker for HIV-1 infectivity, was determined using fluorescence-activated cell sorter (FACS) analysis on Guava easyCyte HT (Millipore Sigma, USA).

### HIV-1 infection of target cells for quantitative assays

Jurkat or SupT1 cells (1 × 10^6^ cells or 2 × 10^6^ cells per well) in 24-well plates were spinoculated with virus stocks (MOI of 0.05 or 0.1) at 480 × *g* for 2 h at 25°C. After the spin, the spinoculated samples were individually transferred onto 1.7 mL microcentrifuge tubes, centrifuged at 300 × *g* for 10 min at 25°C, and the supernatants were carefully removed and discarded. Each cell pellet was then thoroughly resuspended in 1 mL of RPMI complete medium, transferred onto new 24-well cell culture plate, and cultured for 24 h at 37°C. The cells were then harvested by transferring the cell cultures to 1.7-mL microcentrifuge tubes and centrifuging at 1,500 × *g* for 5 min at 25°C. The cell pellets were washed once with PBS and then resuspended in 150 μL of PBS or nuclease-free water, followed by isolation of total DNA using the Quick-DNA miniprep kit (Zymo Research, USA) according to the manufacturer’s instructions.

### Quantitation of reverse transcription products and 2-LTR circles by qPCR

A SYBR green-based qPCR was used to quantify the reverse transcription products, and a TaqMan probe-based qPCR was used to quantify the 2-LTR circles. The qPCRs were assembled in triplicates using 100 ng of sample DNA, 1× iTaq Universal SYBR Green Supermix or 1× iTaq Universal Probe Supermix (Bio-Rad, USA), 300 nM each of forward primer (late RT product: MH531/5′-TGTGTGCCCGTCTGTTGTGT-3′; 2-LTR circle/5′-AACTAGGGAACCCACTGCTTAAG-3′) and reverse primer (late RT product: MH532/5′- GAGTCCTGCGTCGAGAGAGC-3′; 2-LTR circle/5′-TCCACAGATCAAGGATATCTTGTC-3′), and, in the case of probe-based qPCR, 100 nM TaqMan probe (5′-[FAM] ACACTACTTGAAGCACTCAAGGCAAGCTTT-[TAMRA]-3′). The qPCRs were carried out under the following cycling conditions: late RT products- 95°C for 3 min, followed by 39 cycles of amplification and acquisition at 94°C for 15 s, 58°C for 30 s, and 72°C for 30 s; 2-LTR circles- 95°C for 3 min, followed by 39 cycles of amplification and acquisition at 94°C for 15 s, 58°C for 30 s, and 72°C for 30 s. The thermal profile for melting-curve analysis of SYBR green-based qPCR was obtained by holding the sample at 65°C for 31 s, followed by a linear ramp in temperature from 65 to 95°C with a ramp rate of 0.5°C/s and acquisition at 0.5°C intervals. The CFX Manager software (Bio-Rad, USA) was used to determine the copy number of the target DNA in the qPCR samples by plotting the data against the respective standard curve that was generated in parallel using 10-fold serial dilutions of known copy numbers (10^0^ to 10^8^) of pNL43 (for late RT products) or the p2LTR plasmid containing the 2-LTR junction sequence (for 2-LTR circles).

### Quantitation of proviral DNAs by nested PCR

To quantify the copy number of (chromosomally integrated) proviral DNA in HIV-1-infected cells, a nested PCR strategy wherein primers designed to amplify only the junctions of the chromosomal-integrated viral DNA but not any unintegrated viral DNA is used in the first-round end-point PCR, followed by the second round qPCR with primers that amplify viral LTR-specific sequences present in the first round PCR amplicons. The first-round PCR was assembled using 100 ng of sample DNA, 1× GoTaq reaction buffer (Promega, USA), deoxynucleoside triphosphate (dNTP) nucleotide mix containing 200 μM of each nucleotide (Promega, USA), 500 nM each of primers complementary to the target cell chromosomal *Alu* repeat sequence (5′- GCCTCCCAAAGTGCTGGGATTACAG-3) and HIV-1 Gag sequence (5′-GTTCCTGCTATGTCACTTCC-3′), and 1.25 U of GoTaq DNA polymerase (Promega, USA), and the PCR cycling conditions comprised of an initial incubation at 95°C for 5 min, followed by 23 cycles of amplification at 94°C for 30 s, 50°C for 30 s, 72°C for 4 min; and a final extension at 72°C for 10 min. One-tenth volume of the first-round PCR products was used as the template DNA in the second-round qPCR, that also included 1× iTaq Universal Probe Supermix (Bio-Rad, USA), 300 nM each of the viral LTR-specific primers complementary to the LTR-R (5′- TCTGGCTAACTAGGGAACCCA-3′) and the LTR-U5 (5′-CTGACTAAAAGGGTCTGAGG-3′), and 100 nM of TaqMan probe (5′-[6-FAM]-TTAAGCCTCAATAAAGCTTGCCTTGAGTGC-[TAMRA]-3′), and the qPCR was carried out as follows: 95°C for 3 min, 39 cycles of amplification and acquisition at 94°C for 15 s, 58°C for 30 s, and 72°C for 30 s. The CFX Manager software (Bio-Rad, USA) was used to determine the copy number of the target DNA in the qPCR samples by plotting the data against the standard curve that was generated in parallel using 10-fold serial dilutions of known copy numbers (10^0^ to 10^8^) of pNL43.

### Isolation of HIV-1 cytoplasmic PICs and determination of PIC-associated integration activity *in vitro*

The HIV-1 PICs were isolated using a published protocol^74^ with modifications. Briefly, target cells (10 × 10^6^), distributed as 2 × 10^6^ cells per well in 12-well plates, were spinoculated with virus stocks (MOI of 0.05) at 480 × *g* for 2 h at 25°C. The spinoculated samples were individually transferred onto 1.7 mL microcentrifuge tubes, centrifuged at 300 × *g* for 10 min at 25°C, and the supernatants were carefully removed and discarded. Each cell pellet was then thoroughly resuspended in 1 mL of RPMI complete medium, the spinoculated samples corresponding to each virus infection was pooled and distributed as 1 mL aliquots into a new 12-well plate and cultured for 5 h at 37°C. Unless indicated otherwise, all centrifugation steps were carried out at room temperature (∼25°C). The cell samples corresponding to each virus infection was pooled, pelleted by centrifugation at 300 × *g* for 10 min, washed twice with 1 mL of buffer K^−/−^ (20 mM HEPES pH 7.6, 150 mM potassium chloride, 5 mM magnesium chloride), thoroughly resuspended in 0.5 mL of ice-cold buffer K^+/+^ (20 mM HEPES pH 7.6, 150 mM potassium chloride, 5 mM magnesium chloride, 1 mM dithiothreitol [DTT], 20 μg/mL aprotinin, 0.025% [wt/vol] digitonin), incubated on a rocking platform shaker (60 to 80 rocking motions/min) for 10 min at room temperature, centrifuged at 1,500 × *g* for 4 min at 4°C, and the supernatant was collected and centrifuged at 16,000 × *g* for 1 min at 4°C. The resulting supernatant (cytoplasmic PIC) was treated with RNase A (Invitrogen/Thermo Scientific, USA) at a final concentration of 20 μg/mL for 30 min at room temperature, mixed with 60% sucrose (wt/vol) to a final concentration of 7%, and aliquots were flash-frozen in liquid nitrogen and placed in a –80°C freezer for long-term storage. *In vitro* integration assays of cytoplasmic PICs were performed as described previously ^74^. Briefly, the *in vitro* integration reaction was carried out by mixing purified ФX174 target DNA amplicon with 50µL or 100µL of PICs and then incubating the mixture at 37°C for 45 min. The integration reaction was stopped and deproteinized by adding SDS, EDTA, and proteinase K to final concentrations of 0.5%, 8 mM, and 0.5 mg/ml, respectively, and incubating overnight at 56°C. The DNA was purified from the sample by phenol chloroform extraction followed by ethanol precipitation and was used as the template DNA in the first round of the nested PCR. The first-round endpoint PCR, designed to amplify only the integrated viral DNA-target DNA junctions, was carried out in a final volume of 50 µL containing one-tenth volume of the purified DNA from the in vitro integration reaction, 500 nM each of primers targeting the viral LTR (5’- GTGCGCGCTTCAGCAAG-3’) and the target DNA (5’-CACTGACCCTCAGCAATCTTA-3’), 1X GoTaq reaction buffer (Promega, USA), dNTP nucleotide mix containing 200 µM concentrations of each nucleotide (Promega, USA), and 1.25U of GoTaq DNA polymerase (Promega, USA), and under the following thermocycling conditions: initial denaturation at 95°C for 5 min; followed by 23 cycles at 94°C for 30 s, 55°C for 30 s, and 72 °C for 4 min; and a final extension at 72°C for 10 min. The second-round qPCR, designed to amplify only the viral LTR-specific region, was assembled using one-tenth volume of the first-round PCR products as template DNA, 1X iTaq Universal Probe Supermix (Bio-Rad, USA), 300 nM each of the viral LTR-specific primers complementary to the LTR-R (5′-TCTGGCTAACTAGGGAACCCA-3′) and the LTR-U5 (5′- CTGACTAAAAGGGTCTGAGG-3′), and 100 nM of TaqMan probe (5′-[6-FAM]-TTAAGCCTCAATAAAGCTTGCCTTGAGTGC-[TAMRA]-3′), and the qPCR was carried out as follows: 95°C for 3 min, 39 cycles of amplification and acquisition at 94°C for 15 s, 58°C for 30 s, and 72°C for 30 s. The CFX Manager software (Bio-Rad, USA) was used to determine the copy number of the target DNA in the qPCR samples by plotting the data against the standard curve that was generated in parallel using 10-fold serial dilutions of known copy numbers (10^0^ to 10^8^) of pNL43 plasmid. To determine the integration efficiency (i.e., ratio of integrated viral DNA copy numbers to corresponding PIC-associated viral DNA copy numbers), the PIC-associated viral DNA copy numbers were determined. Briefly, 50µL or 100µL of PICs was deproteinized, extracted with phenol-chloroform, and subjected to ethanol precipitation. One-tenth volume of the resulting purified DNA was directly used in qPCR designed to quantify the reverse transcription products. The CFX Manager software (Bio-Rad, USA) was used to determine the copy number of the target DNA in the qPCR samples by plotting the data against the standard curve that was generated in parallel.

### Sequence analysis of junctions of 2-LTR circles from HIV-1-infected target cells

A nested PCR strategy was used to amplify the junction of the 2-LTR circles in HIV-1- infected cells. The first-round end-point PCR was assembled using 500 ng of total DNA from HIV- 1-infected cells, 5× Phusion HF buffer (NEB, USA), a dNTP nucleotide mix containing 200 μM of each nucleotide (Promega), 500 nM each of the primers 2-LTR Forward-M (5′- AGCCTGGGAGCTCTCTGGCTAAC-3′) and 2-LTR Reverse-M (5′-AGCCTTGTGTGTGGTAGATCCAC-3′), and 1 U of Phusion Hot Start Flex polymerase (NEB, USA), and the PCR was carried out using cycling conditions of 98°C for 2 min, 30 cycles of amplification at 98°C for 10 s, 71°C for 1 min, and 72°C for 15 s, and a final extension at 72°C for 10 min. The second-round end-point PCR was assembled using 1/10^th^ volume of the first-round PCR products as template DNA, along with 5× Colorless GoTaq reaction buffer, 500 nM each of the primers MH535 (5′-AACTAGGGAACCCACTGCTTAAG-3′) and MH536 (5′-TCCACAGATCAAGGATATCTTGTC-3′), and 1.25 U of GoTaq DNA polymerase, and the PCR was carried out under the following cycling conditions: 95°C for 5 min, 25 cycles of amplification at 95°C for 15 s, 58°C for 1 min, and 72°C for 30 s, and a final extension at 72°C for 10 min. The resulting PCR amplicon from the second-round PCR was gel-purified and ligated to the pGEMT-Easy vector (Promega, USA), as per the manufacturer-recommended protocol (Promega, USA). Diagnostic restriction digestion was used to verify the presence of 2-LTR junction DNA insert in the recombinant plasmid DNA isolated from transformed bacterial colonies selected on X-Gal (5- bromo-4-chloro-3-indolyl-β-d-galactopyranoside)/IPTG (isopropyl-β-d-thiogalactopyranoside) indicator plates. The sequence of the 2-LTR junction DNA inserts was determined by Sanger DNA sequencing using primers flanking the cloning site, and multiple alignment was used to analyze the sequence data.

## ACKNOWLEDGEMENTS

This work was supported by the National Institutes of Health grants R01 AI136740, R01 DA 042348, R56 AI122960, R24 DA036420, and U54 MD007586 to C.D. and a Research Centers in Minority Institutions (RCMI) grant U54MD007586 to J.P. This work is also supported in part by the Meharry Translational Research Center (MeTRC) grant U54MD007593 and Tennessee CFAR grant P30 AI110527 from the National Institutes of Health to C.D.

## Notes

### Competing Interest Statement

The authors have declared no competing interest.

