## Supplemental File for "HIV-1 Mutants that Escape the Cytotoxic T-Lymphocytes are Defective in Viral DNA Integration"

#### SUPPLEMENTARY FIGURE LEGENDS

**Figure S1.** Electrostatic potential surfaces of the wildtype and three mutant hexamers. The CA hexamers are shown in molecular surface representations. The magnitude of the electrostatic potential is represented by the colors on the molecular surface. The regions of mutated residues are highlighted, indicating the electrostatic potential changes after residue substitutions.

**Figure S2.** Ion occupancies of CA hexamers. **(A)** Ion occupancies of four CA hexamer systems from CIONIZE. Occupancy of Na and Cl ion is in yellow and cyan respectively and the isovalue of the density map shown is 0.01 for both ions. **(B)** Relative chloride ion occupancy differences of three mutant hexamers, with respect to a selected reference system. The $Cl^-$ ion occupancies of the RK hexamer with reference to WT hexamer, RKLM hexamer to RK hexamer, and RKLMSA hexamer to RKLM hexamer, are shown from left to right. The occupancies in blue represent chloride ion occupancy present in the test system but not in the reference system. Occupancies in lime representing ions present in the reference system but not in the test system.

**Figure S3.** Effect of KK10-linked CTL escape mutation R264K on the integration activity in vitro of cytoplasmic PICs**.** Integration activity in vitro and the viral DNA content of the WT and RK cytoplasmic PICs isolated from SupT1 cells spinoculated with the respective viruses for 2 hours at 25°C and cultured for 5 hours at 37°C were determined as described in Materials and Methods. **(A)** Integration activity of the cytoplasmic PICs, i.e., the copy number of viral DNAs that were integrated into the target DNA during the in vitro integration assay, as measured by qPCR and calculated using a standard graph of known copy numbers of HIV-1 molecular clone. **(B)** Percentage integration activity plotted relative to WT. **(C)** Copy number of viral DNA content in the cytoplasmic PICs as measured by qPCR and calculated using a standard graph of known copy numbers of HIV-1 molecular clone. **(D)** Percentage viral DNA content plotted relative to WT cytoplasmic PICs. **(E)** Ratio of the copy number of integrated viral DNAs to the corresponding PIC-associated viral DNAs plotted as percentage relative to WT. Data shown are representative of three independent experiments, with error bars representing the SEMs.

**Figure S4.** Assessment of the effect of KK10-linked CTL escape mutations on the integrity of the termini of the reverse transcription products (viral DNAs). Junctions of the 2-LTR circles present in the total DNA isolated from Jurkat cells spinoculated with the WT or mutant viruses for 2 hours at 25°C and cultured for 24 hours at 37°C were amplified by a nested PCR strategy, cloned into a plasmid vector, and the DNA sequences determined by Sanger sequencing were aligned and analyzed as described in Materials and Methods. Shown are the 2-LTR circle junction sequences from cells inoculated with **(A)** WT virus, **(B)** R264K virus, and **(C)** R264K/L268M/S173A virus. The 3'U5 sequence is in blue, the 5'U3 sequence is in red, the unprocessed terminal dinucleotides (GT and AC) are in black, the deletions within the 3'U5 or 5'U3 region are indicated by dashes, and the DNA insertions are in magenta. The insertion sequences highlighted in yellow or blue correspond to the PPT or PBS regions, respectively.

### Figure S1

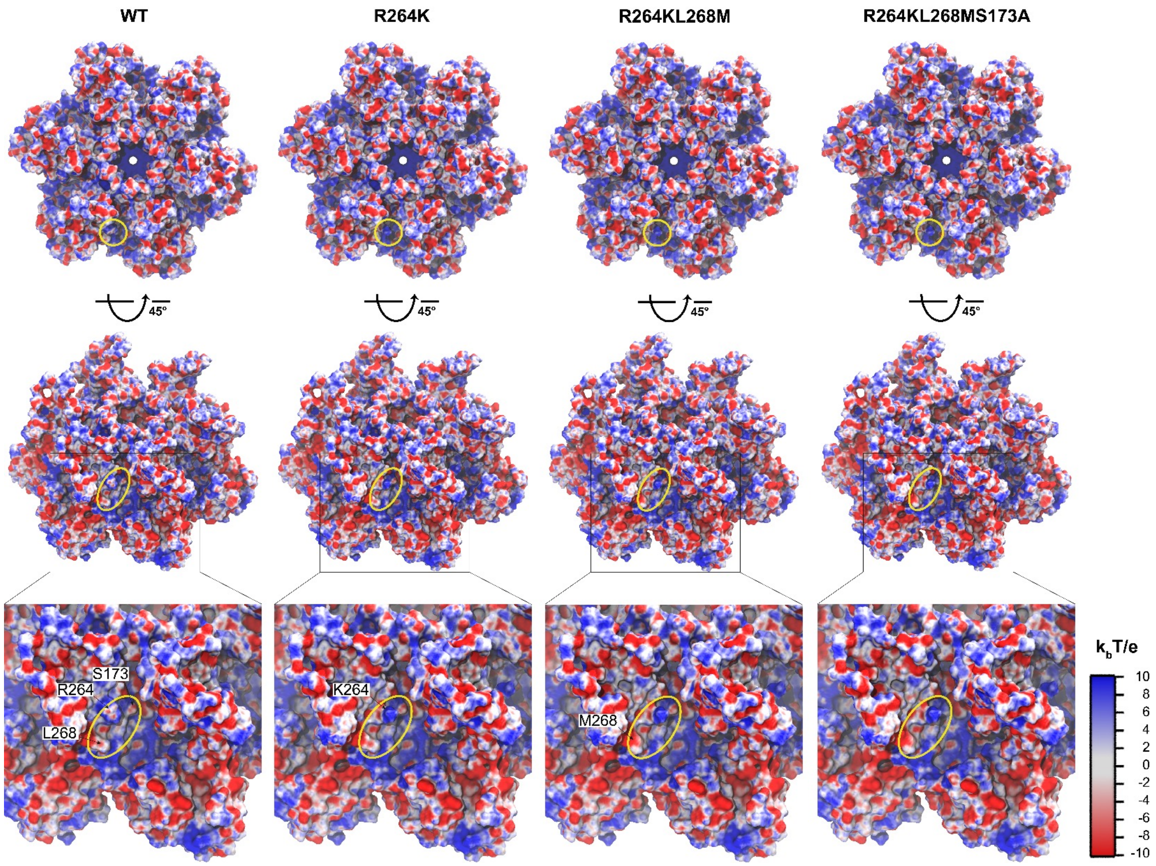

WT
R264K
R264KL268M
R264KL268MS173A
45°
45°
45°
45°
S173
R264
L268
K264
M268
kbT/e
10
8
6
4
2
0
-2
-4
-6
-8
-10

### Figure S2

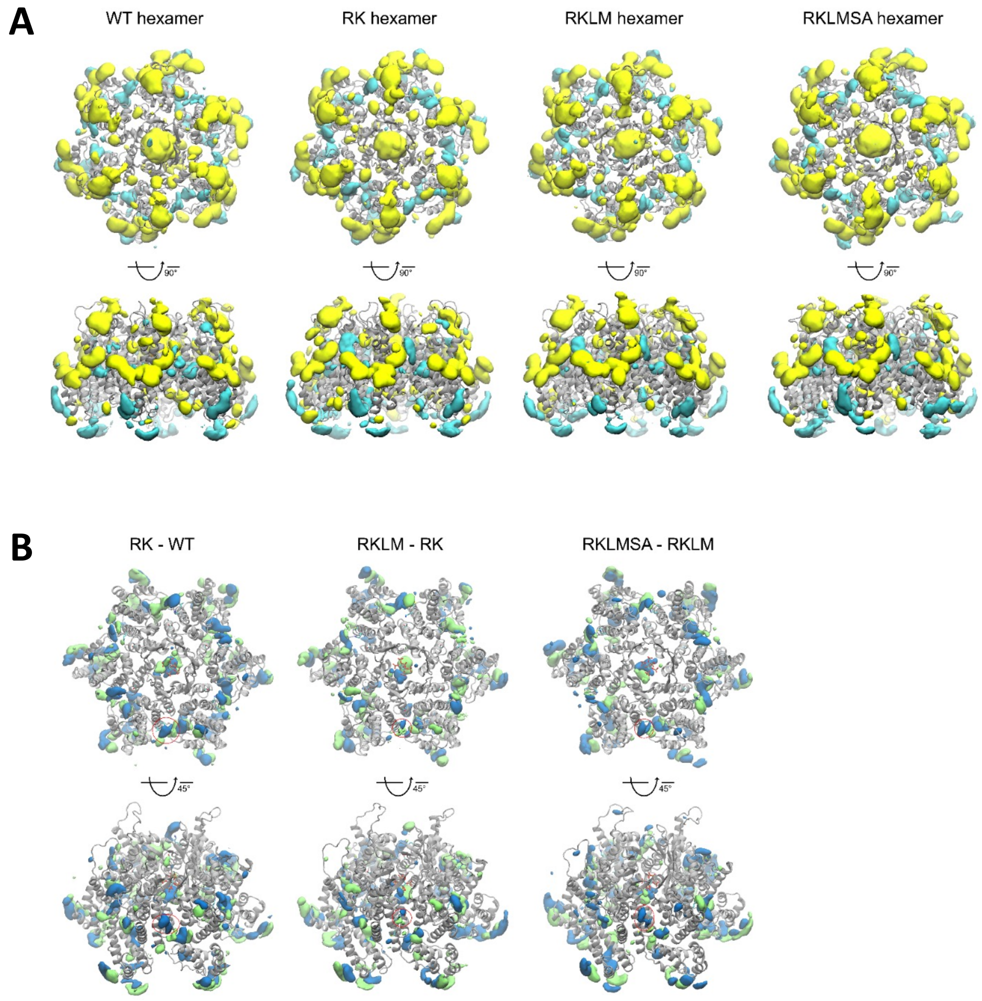
A
WT hexamer
RK hexamer
RKLM hexamer
RKLMSA hexamer
90°
B
RK - WT
RKLM - RK
RKLMSA - RKLM
45°

#### Figure S3

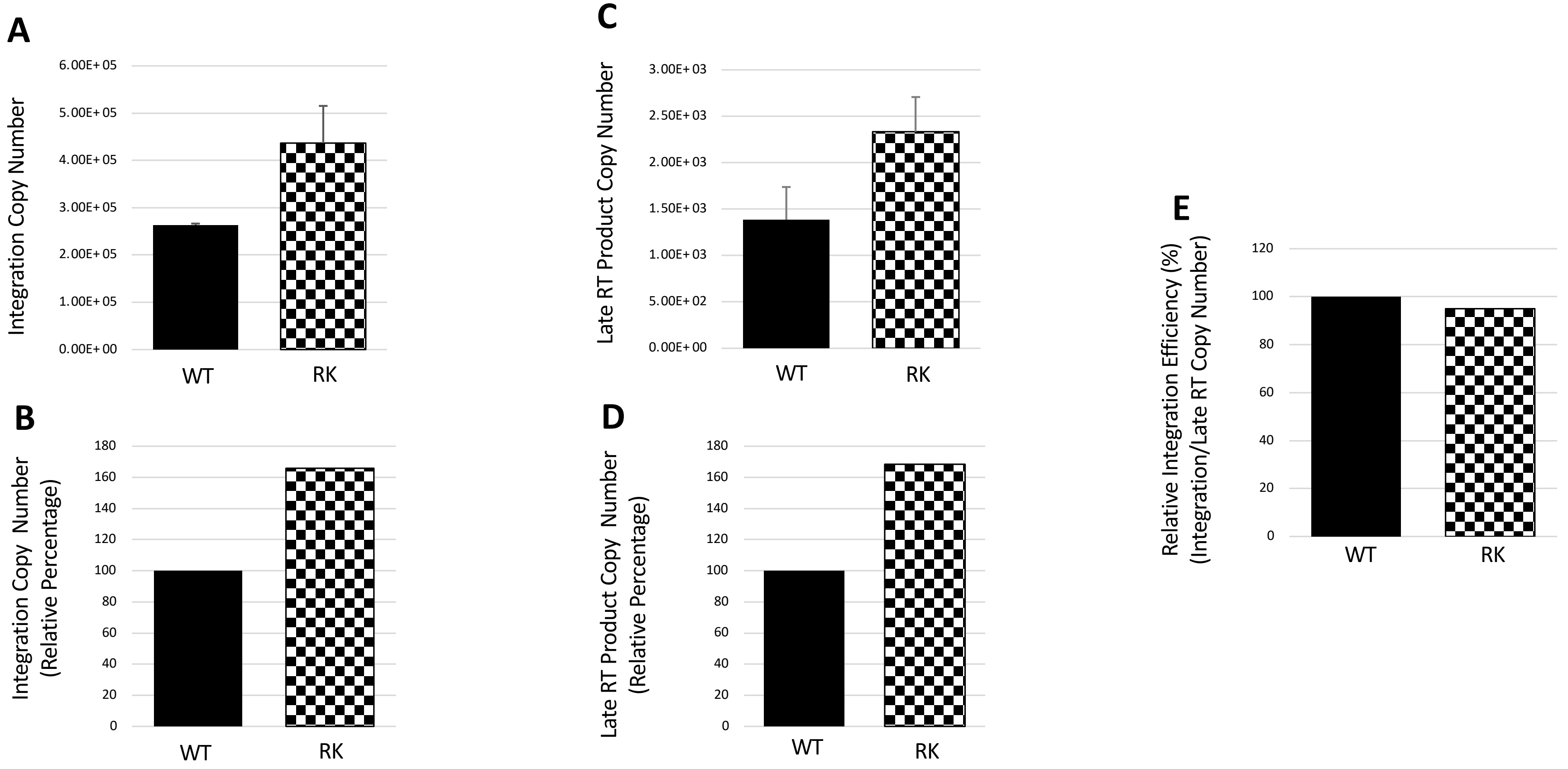

A
Integration Copy Number
6.00E+05
5.00E+05
4.00E+05
3.00E+05
2.00E+05
1.00E+05
0.00E+00
WT
RK
B
Integration Copy Number
(Relative Percentage)
180
160
140
120
100
80
60
40
20
0
WT
RK
C
Late RT Product Copy Number
3.00E+03
2.50E+03
2.00E+03
1.50E+03
1.00E+03
5.00E+02
0.00E+00
WT
RK
D
Late RT Product Copy Number
(Relative Percentage)
180
160
140
120
100
80
60
40
20
0
WT
RK
E
Relative Integration Efficiency (%)
(Integration/Late RT Copy Number)
120
100
80
60
40
20
0
WT
RK

### Figure S4

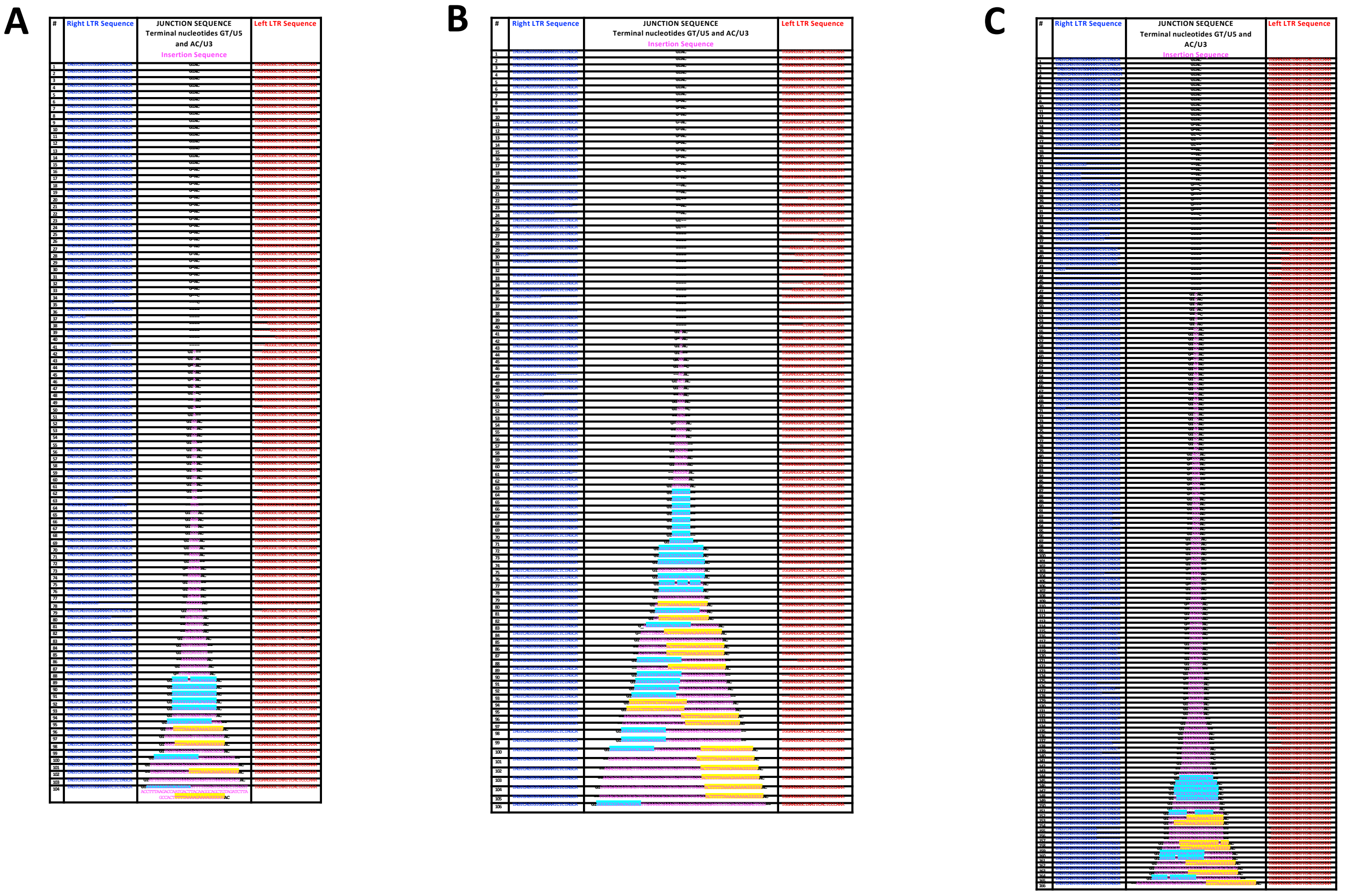
A
B
C
#
Right LTR Sequence
JUNCTION SEQUENCE
Terminal nucleotides GT/U5 and AC/U3
Insertion Sequence
Left LTR Sequence
